# The microprotein Dafcin resembles influenza HA fusion peptide and regulates the size of storage lysosomes in the germline

**DOI:** 10.64898/2026.07.08.737289

**Authors:** Kevin G. Nyberg, Ryder Easterlin, Cooper W. P. Stringer, Hamdi K. Küçükengin, Manny Joseph Widuch, Kyung Je Lee, Silpol Dhiantravan, Megan A. Wong, Richard W. Carthew

## Abstract

Microproteins translated from short open reading frames are increasingly understood to play important roles in cell biology and development. Here, we describe a microprotein in *Drosophila* that is expressed in ovarian follicle cells which surround the developing oocyte. The Dafcin microprotein is predicted to form an amphipathic alpha-helix, a structure known to interact with lipid bilayers. Dafcin’s structure most resembles the influenza HA fusion peptide, which induces negative curvature of endosomal membranes. Dafcin tagged with GFP localizes to the Golgi and is ultimately secreted from the follicle cells. Remarkably, this occurs without the microprotein having a secretory signal sequence. The protein is taken up into the oocyte by endocytosis, localizing to the inner face of storage lysosomes called yolk granules. Mutant analysis shows that Dafcin is required to limit the size of yolk granules. This may occur by inducing negative membrane curvature like HA peptide. In support, liposomes formed in vitro with both Dafcin and HA peptides are smaller in size.

## Introduction

It has become apparent that large numbers of genes annotated as long noncoding RNA (lncRNA) genes actually encode proteins of 100 amino acids or less called microproteins^1–5^. Historically, gene annotation used an open reading frame (ORF) length threshold of 100 codons to classify genes as protein-coding^6^. However, evidence from proteomics and Ribo-Seq experiments point to widespread translation of ORFs smaller than 100 codons, termed short ORFs (sORFs), from annotated lncRNAs as well as 5’ UTRs. These sORFs number in the hundreds to thousands for bacteria^7^ and eukaryotes alike, including humans^1–3,5^. The functional importance of some microproteins is supported by studies of individual genes and by forward genetic screens. In *Drosophila*, for example, the classic Polished rice/Tarsal-less microproteins are 11-32 amino acids in length and function to regulate the activation of the transcription factor Shavenbaby^8,9^. Humanin, a 21 or 24-amino acid microprotein encoded from the mitochondrial 16S rRNA region, has been shown to protect cells against apoptosis driven by Alzheimer’s disease genes^10–12^. Moreover, some microproteins maintain sequence conservation across evolutionary time, hinting at functional significance^1,13^. However, most microproteins have unknown functions and comprise an extensive and intriguing ‘dark proteome’.

The structures of microproteins, both predicted and solved, display a high degree of diversity^3,14^. In most cases, these structures do not resemble those adopted by secreted peptides, which are typically processed by proteolysis to generate their active forms. For example, the class of secreted peptides that adopt an amphipathic alpha-helix conformation include anti-microbial peptides (AMPs) and cell penetrating peptides (CPPs), the latter of which have been engineered for therapeutic purposes^15–17^. AMPs and CPPs are composed of an alpha-helix with opposing hydrophobic and hydrophilic surfaces. Microproteins with amphipathic helical structures, however, are rare. One example is SpoVM from *Bacillus subtilis*, which folds into a kinked helix with a hydrophobic surface and a positively charged polar surface^18,19^. The microprotein binds to lipid membranes via its hydrophobic surface, with preference for membranes with positive curvature^20^. Once bound, SpoVM recruits the ATPase SpoIVA to the membrane via binding to its hydrophilic face^21^. This interaction is required for *B. subtilis* to form spores under adverse environmental conditions. AzuC of *Escherichia coli* is another amphipathic microprotein that also binds the cell membrane and recruits a larger protein^22^. In this case, it binds glycerol 3-phosphate dehydrogenase and stimulates its enzymatic activity.

Amphipathic helices also are found within many proteins not classified as microproteins. The helices allow proteins to bind the lipid surface of cell organelles where they can perform a variety of functions, including membrane deformation and membrane curvature-sensing^23,24^. A classic example is Hemagglutinin (HA) glycoprotein, which coats the membrane surface of the influenza virus^25,26^. The HA2 subunit contains a 23-amino acid domain near its N-terminus, which folds into a kinked amphipathic helix called the HA fusion peptide^27,28^. Once a virus particle enters a cell by endocytosis, the drop in endosome pH triggers a conformational change in HA, and the HA fusion peptide inserts into the inner monolayer of the endosome membrane^29,30^. This anchors HA to the endosome membrane. After a series of further conformational changes, HA brings the virus membrane into close proximity to the endosome membrane with the inserted fusion peptide^29^. The two monolayers fuse to form a stalk, which resolves to an open pore through which the virus particle can pass, entering the cytosol. The fusion peptide lowers the barrier to stalk formation by promoting strong negative curvature of the stalk^31^. Thus, the fusion peptide is essential for membrane fusion.

Here, we describe a 21-amino acid microprotein expressed in the fruit fly *Drosophila melanogaster* predicted to form an amphipathic alpha-helix structurally similar to HA fusion peptide. The Dafcin microprotein is synthesized in follicle cells of the ovary and enters the secretory pathway without the presence of an ER signal sequence^32^. Dafcin is taken up by oocyte cells via endocytosis and is retained in these organelles as they mature into storage lysosomes containing yolk proteins. It occupies a cortical layer adjacent to the inner monolayer of the membrane. Dafcin is required to limit the size of the storage lysosomes, and this may occur via inducing membrane curvature. Like HA fusion peptide, Dafcin co-incubated with lipids in vitro induces them to form smaller liposomes. Thus, Dafcin functions to regulate storage granule maturation using a biochemical activity reminiscent of influenza fusion peptide.

## Results

### Expression of a conserved microprotein in the ovary

The *CR43834* gene is one of approximately 2,500 genes annotated as lncRNA genes in the *Drosophila melanogaster* genome (BDGP r6.65). *CR43834* RNA is detectable in ovary RNA-Seq datasets from several species of *Drosophila* (*melanogaster*, *simulans*, and *yakuba*), indicating that the developmental regulation of *CR43834* may be conserved. Using RNA single-molecule fluorescent in situ hybridization (smFISH) in *D. melanogaster* ovaries, *CR43834* RNA was detected in the somatic follicle cells surrounding the oocyte from stage 10B through to the end of oogenesis (Fig. 1A).

**Figure 1.**
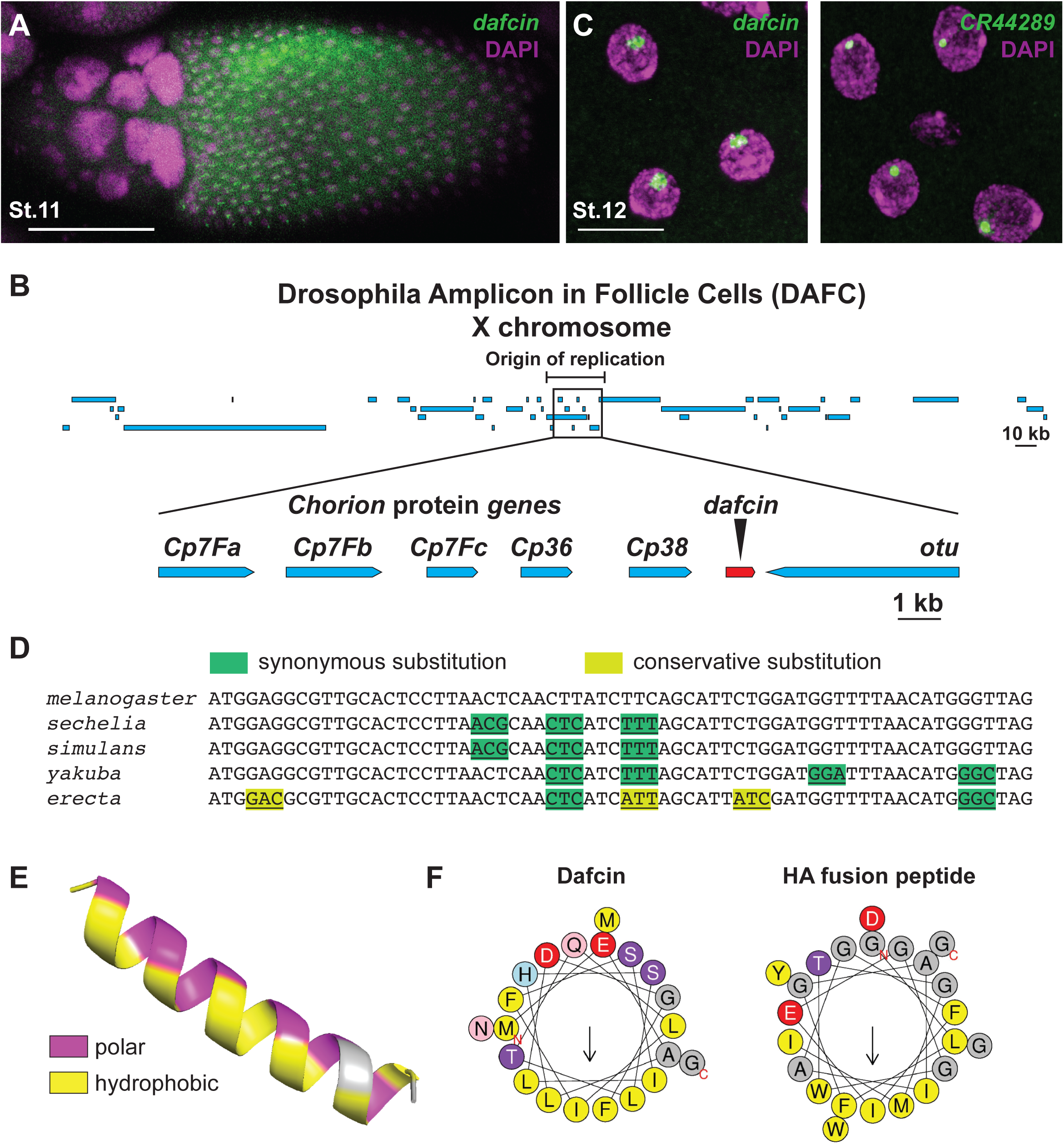
The *dafcin* gene, located in a high-copy number genomic region, produces an amphipathic alpha-helical microprotein in ovarian follicle cells. **(A)** smFISH staining of *dafcin* RNA (green) in a stage 11 egg chamber. The large nuclei at the anterior (left) end belong to germline nurse cells. The smaller nuclei belonging to the somatic follicle cells cover the posterior domain of the egg chamber. Nuclei are counterstained with DAPI. Scale bar represents 100 μm. **(B)** The *dafcin* gene is located in a DAFC region on the X chromosome, adjacent to a cluster of chorion protein genes^37^. The origin of replication of the DAFC region maps to a 25-kb region around the cluster^36^. **(C)** High magnification of stage 12 follicle cells stained by smFISH for *dafcin* (left) or *CR44289* RNA (right). The single fluorescent spots localized within each nucleus correspond to smFISH probes hybridized to nascent RNAs at the site of transcription. The size of *dafcin* RNA transcription spots is much larger than the size of spots corresponding to *CR44289*, a gene located in a non-amplified region of the genome. Scale bar represents 10 μm. **(D)** The 63-nucleotide *dafcin* sORF in species of the *melanogaster* subgroup have either synonymous or conservative amino acid substitutions, consistent with purifying selection in the coding region. **(E)** Using AlphaFold 3 and PyMOL, Dafcin is predicted to form an amphipathic alpha-helix^64,65^. Hydrophobic residues are depicted in yellow, and polar (i.e., hydrophilic) residues are depicted in magenta. **(F)** Helical wheel plots illustrating the properties of the Dafcin and HA fusion peptide alpha helices^66^. Hydrophobic residues are depicted in yellow, acidic residues in red, basic residues in blue, polar uncharged residues in purple, and the small aliphatic residues glycine and alanine in gray. The predicted Dafcin microprotein shares striking physico-chemical similarities with the influenza HA fusion peptide, including similar hydrophobicities, hydrophobic moments, and -2 net charge.

By stage 10B, the follicle cells surrounding the oocyte have undergone three rounds of whole genome endoreplication^33,34^. Localized endoreplication then continues in six regions of the genome, ultimately producing a higher copy number of genes located within these amplified regions relative to the rest of the genome^35–38^. Known as Drosophila Amplicons in Follicle Cells, or DAFCs, several of the amplified regions contain clusters of chorion genes, and DAFCs are thought to have evolved to facilitate the synthesis of large quantities of chorion proteins in a short developmental timeframe^35,37^. The *CR43834* gene is located within a DAFC on the X chromosome near its origin of amplification (Fig. 1B)^36^. Indeed, *CR43834* RNA was detected by smFISH within each follicle cell nucleus in an unusually large fluorescent focus, presumably corresponding to nascent transcripts made from the amplified gene (Fig. 1C). Due to the unique location of *CR43834* within a DAFC, we decided to rename the gene as *dafcin* (*dfcn*).

Although *dafcin* is annotated as a lncRNA-coding gene, PhyloCSF analysis detected a sORF within the *dafcin* transcript that is conserved among several *Drosophila* species within the *melanogaster* subgroup (Fig. 1D)^39^. The sORF has an invariant length of 63 nucleotides among these species, and all predicted codon substitutions between species are either synonymous or conservative. This strongly suggests that the sORF encodes a conserved 21-amino acid microprotein. The encoded microprotein is predicted to form an amphipathic alpha-helix, with hydrophobic residues localized to one face of the alpha-helix, and hydrophilic (i.e., polar) residues localized to the opposing face (Fig. 1E). The hydrophilic face is predicted to include two negatively charged residues.

Amphipathic peptides often interact with lipid bilayers and perform diverse functions, including cell penetration and viral-host membrane fusion^15,16,40,41^. Typically, the polar face of such peptides is positively charged in order to facilitate their association with phospholipids^23,42^. However, the Dafcin peptide has a negatively charged polar face, which is rare among amphipathic helices. The HA2 protein coating influenza virus particles contains one such negatively charged amphipathic helix (Fig. 1F)^27,29,43^. This HA fusion peptide is essential for HA protein-triggered fusion of the virus and host membranes and subsequent virus penetration into the cytosol^44^. Both Dafcin and HA fusion peptides have two acidic residues on their hydrophilic surfaces. Strikingly, they also share an unusually strong hydrophobic moment (0.409 and 0.447, respectively) and bulky hydrophobic face (0.706 and 0.732 mean hydrophobicity, respectively) when compared to other amphipathic peptides.

To test for translation of the conserved sORF from *dafcin* RNA, we used CRISPR/Cas9 homology directed repair (HDR) to precisely edit the endogenous *dafcin* gene, inserting superfolder GFP at the carboxyl-terminus of the sORF (Fig 2A). Dafcin-GFP fusion protein was detected in late-stage follicle cells surrounding the oocyte starting at stage 10B, consistent with smFISH analysis of *dafcin* RNA in the ovary (Fig 2B). The abundance of the Dafcin-GFP fusion protein was highly variable from cell-to-cell in the follicle epithelium at stage 10B, though this variation was not patterned along any of the body axes. The heterogeneity typically disappeared at later stages of oogenesis, and expression became more uniform (Fig. 2B). The coding potential of the *Dafcin* sORF was further supported by detection of Dafcin-GFP fusion protein when expressed in *Drosophila* S2 cells (Fig. 2C). Taken together, we provide compelling evidence that a Dafcin microprotein is expressed in vivo.

**Figure 2.**
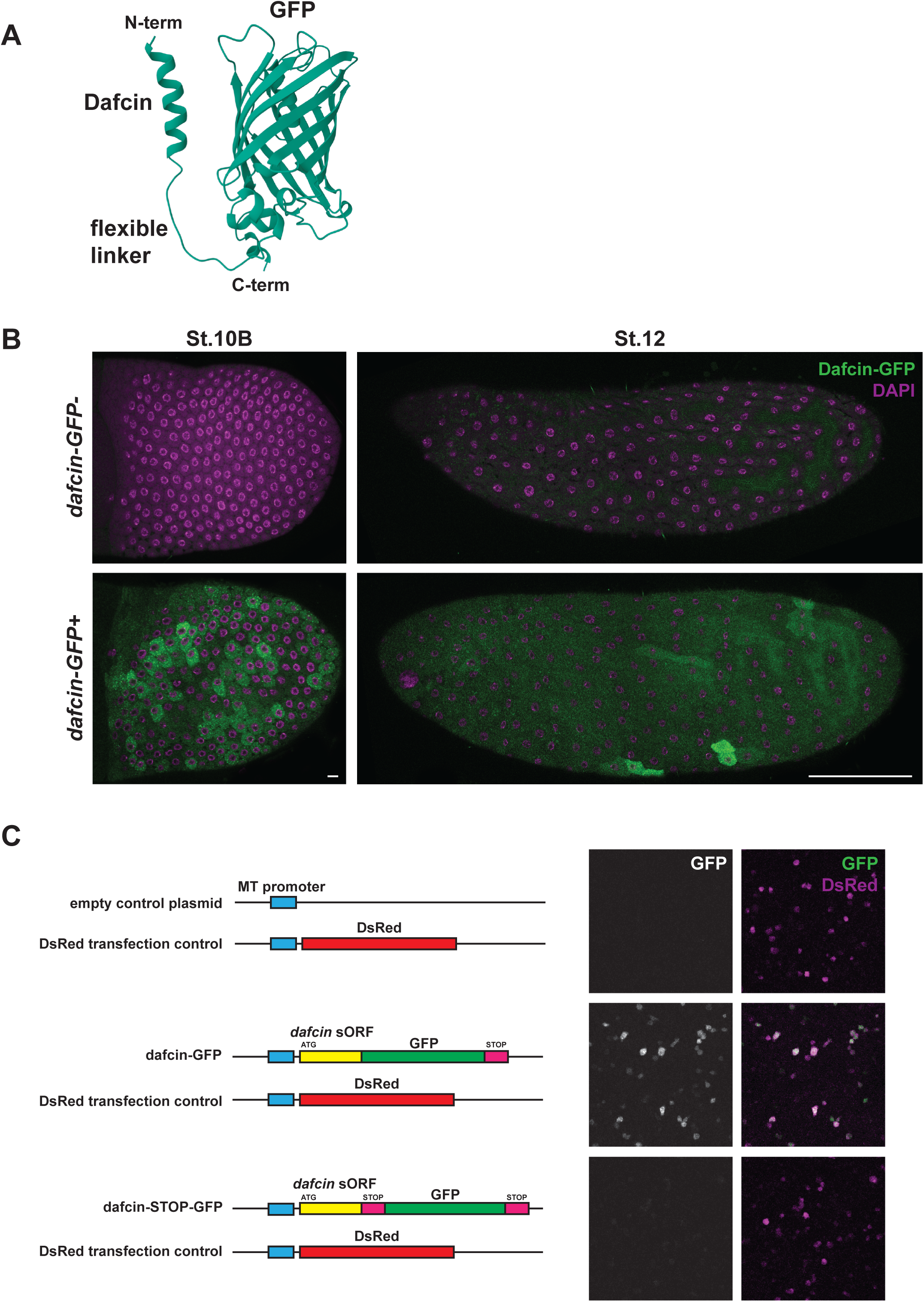
The *dafcin* sORF is translated into protein. **(A)** Predicted structure of the Dafcin-GFP fusion protein produced from the endogenous *dafcin* gene after in-frame insertion of GFP sequences. AlphaFold 3 was used to generate the structure^65^. **(B)** Stages 10B and 12 egg chambers from females carrying the endogenous *dafcin* gene (top panels) or *dafcin-GFP* gene (bottom panels). Dafcin-GFP fluorescence (green) was detected in follicle cells surrounding the oocyte starting at Stage10B. Scale bars represent 10 μm (Stage 10B) and 100 μm (Stage 12). **(C)** Dafcin-GFP expression as detected by green fluorescence after transient transfection of *Drosophila* S2 cells. Expression plasmids were co-transfected with a control plasmid carrying a DsRed marker gene to identify successfully transfected cells. Top, the expression plasmid not carrying any gene. Middle, the expression plasmid carrying GFP sequence in frame with the *dafcin* 63-nt sORF. Bottom, the expression plasmid carrying GFP out of frame with the *dafcin* sORF.

### Unconventional secretion and trafficking of Dafcin-GFP in the ovary

We examined the subcellular localization of Dafcin-GFP within follicle cells. The protein was dispersed throughout the cytoplasm, as well as enriched near the apical cell membrane of stage 10B follicle cells (Fig. 3A). It was also strongly enriched in punctae located throughout the cytoplasm. Some of these punctae co-localized with both cis- and trans-Golgi markers, indicating that Dafcin-GFP localizes to the Golgi (Fig. 3B).

**Figure 3.**
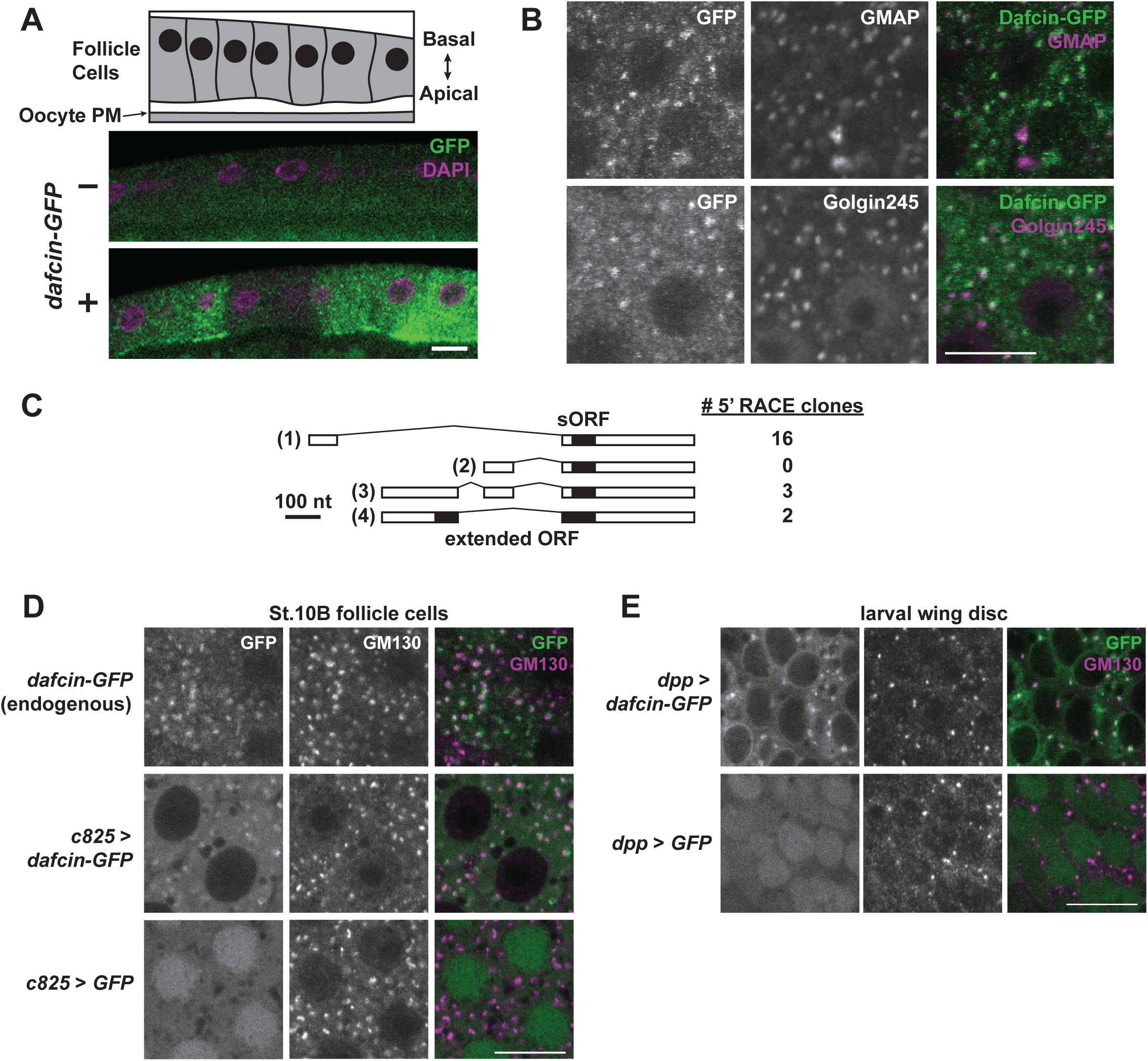
Dafcin-GFP is enriched in the apical domain of follicle cells and localizes to the Golgi. **(A)** Top, schematic of follicle cell epithelium and oocyte plasma membrane (PM). Bottom, Dafcin-GFP fluorescence is localized throughout the follicle cell cytoplasm and is apically enriched in Stage 10B follicle cells. **(B)** Stage 10B follicle cells expressing Dafcin-GFP and co-stained for GMAP (cis-Golgi) and Golgin245 (trans-Golgi) proteins. Images are in the plane of the epithelium. **(C)** 5’ RACE analysis of ovary mRNA probing for *dafcin* transcripts. Three alternate splice isoforms (isoforms 1, 3, and 4) were cloned and characterized, and numbers indicating their relative abundance are to the right. While isoform 2 was not cloned by 5’ RACE, its existence is supported by multiple RNA-Seq datasets from ovaries. The great majority of clones were derived from isoforms 1 and 3 containing the 63-nt sORF (19/21 clones). This result is also consistent with count data from multiple ovary RNA-Seq experiments. **(D)** Stage 10B follicle cells stained for GFP fluorescence (green) and the cis-Golgi marker GM130 (purple). Top, cells expressing endogenous *dafcin-GFP*. Middle, cells expressing *UAS-dafcin-GFP* driven by *c825-Gal4*. Bottom, cells expressing *UAS-GFP* driven by *c825-Gal4*. **(E)** Third-instar larval wing disc cells stained for GFP fluorescence (green) and GM130 (purple). Top, cells expressing *UAS-dafcin-GFP* driven by *dpp-Gal4*. Bottom, cells expressing *UAS-GFP* driven by *dpp-Gal4*. Scale bars in all panels represent 10 μm.

Newly translated proteins destined for the Golgi contain a signal peptide that enables their translocation into the ER for trafficking to the Golgi^32^. The 21-amino acid Dafcin microprotein does not contain a recognizable signal sequence (Supp. Fig. 1). These are typically stretches of hydrophobic residues that fold into an alpha-helix preceded by basic residues. In contrast, the Dafcin helix is amphipathic and acidic in charge. We first considered the possibility that the annotated Dafcin mRNA structure was incorrect, and a larger protein was being translated with a signal sequence. Extensive 5’ RACE was performed on ovary RNA, and virtually all characterized transcripts derived from *dafcin* contained the 63-nucleotide sORF (Fig. 3C). However, there were a few transcripts that appeared to be splice variants that extended the sORF to an upstream exon (isoform 4, Fig. 3C). Indeed, this extended ORF was strongly predicted by SignalP 6.0 to form a signal peptide for ER translocation (Supp. Fig. 1). However, the rarity of these transcripts from the ovary was at odds with the Golgi localization of bulk Dafcin-GFP protein.

To test whether the 21-amino acid Dafcin microprotein was sufficient to traffic GFP to the Golgi, we constructed a Dafcin-GFP transgene under the control of the UAS promoter, using the isoform 2 cDNA of *dafcin*, which is the most abundant isoform in RNA-Seq data (Fig. 3C). This transgene only encodes the 63-nucleotide sORF. When expressed in follicle cells, transgenic Dafcin-GFP was localized to the Golgi, recapitulating the localization observed with endogenous Dafcin-GFP (Fig. 3D). When the transgene was expressed in larval imaginal wing disc cells, Dafcin-GFP was also localized to the Golgi, indicating that follicle cells did not have a unique mechanism for its Golgi localization (Fig. 3E). Moreover, expression of the 63-nucleotide sORF fused to GFP in *Drosophila* S2 cells also showed localization of the fusion protein to their Golgi (data not shown).

Secretion of proteins from cells generally requires their transit through the Golgi. Since Dafcin-GFP was Golgi-localized, we asked whether Dafcin was being secreted from the follicle cells. Apical enrichment of Dafcin-GFP in follicle cells suggested a potential route of secretion, and indeed, we detected Dafcin-GFP in oocytes that were surrounded by Dafcin-expressing follicle cells (Fig. 4A). The oocyte-localized Dafcin-GFP took the form of bright punctae, as well as slightly larger fluorescent rings, which suggested potential localization to cytoplasmic organelles. Transgenic Dafcin-GFP specifically expressed in follicle cells also showed similar oocyte localization (Fig. 4B).

**Figure 4.**
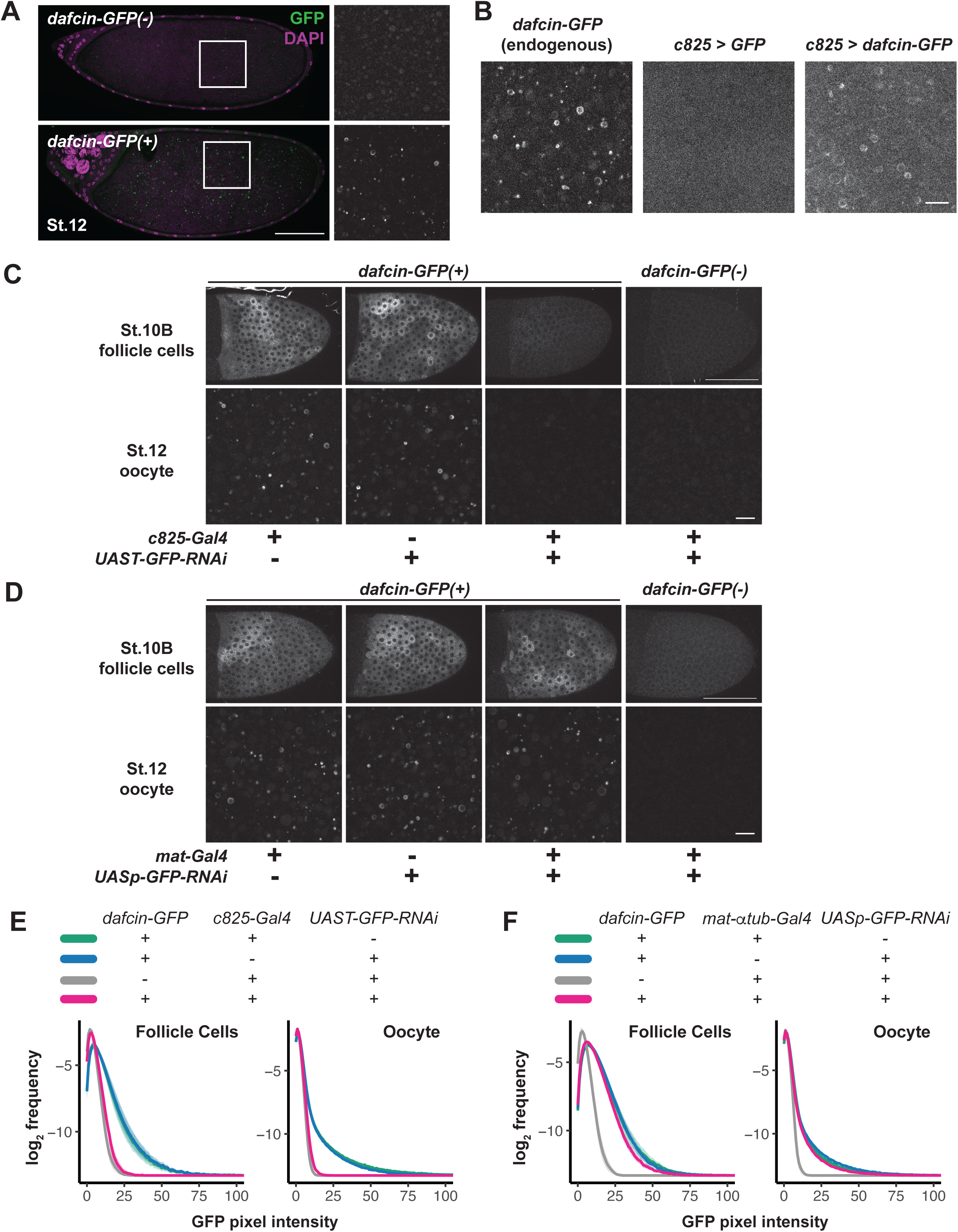
Dafcin-GFP traffics from the follicle cells into the oocyte. **(A)** Mid-sagittal green fluorescence images of stage 12 egg chambers from females carrying the endogenous *dafcin* gene (top panels) or *dafcin-GFP* gene (bottom panels). Anterior is to the left and scale bar represents 100 μm. Right panels show a magnified view of oocyte cytoplasm highlighted in the left panels. **(B)** Magnified images of cytoplasm in stage 12 oocytes. Scale bar represents 10 μm. Left, an oocyte surrounded by follicle cells expressing endogenous *dafcin-GFP*. Middle, an oocyte surrounded by follicle cells expressing *UAS-GFP* driven by *c825-Gal4*. Right, an oocyte surrounded by follicle cells expressing *UAS-dafcin-GFP* driven by *c825-Gal4*. The higher fluorescence backgrounds in the right panels are due to bleed-through from strong GFP or Dafcin-GFP fluorescence in the follicle cells. **(C, D)** RNAi knockdown of *dafcin-GFP* expression using an inducible UAS-RNAi transgene driven by either *c825-Gal4* **(C)** or *maternal α-tubulin-Gal4* (*mat-Gal4*) **(D)**. *c825-Gal4* is specifically expressed in follicle cells whereas *mat-Gal4* is specifically expressed in nurse cells. Top panels show GFP fluorescence in stage 10B follicle cells and bottom panels show GFP in stage 12 oocytes. Scale bars represent 100 μm in St.10B follicles and 10 μm in St.12 oocytes. Samples are taken from either females carrying the endogenous *dafcin* gene or *dafcin-GFP* gene, as indicated. **(E, F)** Quantification of the RNAi knockdown experiments. Shown are frequency distributions of pixel-wise GFP fluorescence intensity in follicle cell and oocyte images. Moving line averages are in solid lines, and standard deviations are in shaded areas. Genotypes are as indicated for RNAi driven by *c825-Gal4* **(E)** and *mat-Gal4* **(F)**. Note that these distributions correspond to data collected from images in (C) and (D), as well as several other replicates.

To fully rule out the possibility that low-level germline expression of *dafcin* accounted for the oocyte signal, we triggered RNAi knockdown of *dafcin-GFP* specifically in follicle cells by targeting *GFP* and observed near-complete absence of Dafcin-GFP protein in the oocyte (Fig. 4C, 4E). RNAi knockdown of *GFP* specifically in the germline had almost no effect on Dafcin-GFP accumulation in the oocyte (Fig. 4D, 4F). To ensure that *GFP* knockdown was generally possible in the germline, we used the same *GFP* RNAi construct to knockdown *oskar-GFP*, which is known to be expressed in the germline nurse cells (Supp. Fig. 2). In sum, these results indicate that the Dafcin microprotein is sufficient to traffic GFP from the follicle cells to the oocyte, even without a classic N-terminal signal sequence.

### Endocytosis of Dafcin-GFP into oocyte yolk granules

Dafcin-GFP was localized within the oocyte to structures resembling cytoplasmic organelles. These organelles possibly originated by endocytosis, thus explaining how secreted Dafcin-GFP entered the oocyte. To test if Dafcin-GFP entered the oocyte via endocytosis, we eliminated the Yolkless (Yl) receptor via mutation of the *yl* gene. Yl is a low-density lipoprotein receptor responsible for the massive uptake of vitellogenins (yolk proteins) by endocytosis into the oocyte^45^. In *yl*^13^ mutant ovaries, endocytosis of yolk proteins, which autofluoresce, was effectively abolished, resulting in the complete absence of yolk proteins in the oocyte (Fig. 5A)^46^. When *dafcin-GFP* was expressed in *yl*^13^ mutant ovaries, there was a dramatic reduction of Dafcin-GFP protein found in the oocyte, though some small punctae remained (Fig. 5A, 5B). Instead, Dafcin-GFP was found to accumulate in the extracellular space between the follicle cells and the oocyte in a subset of egg chambers, with enrichment at both oocyte and follicle cell membranes (Fig. 5C). This observation is consistent with an abnormal buildup of Dafcin-GFP in the extracellular space due to drastically reduced endocytosis by the oocyte.

**Figure 5.**
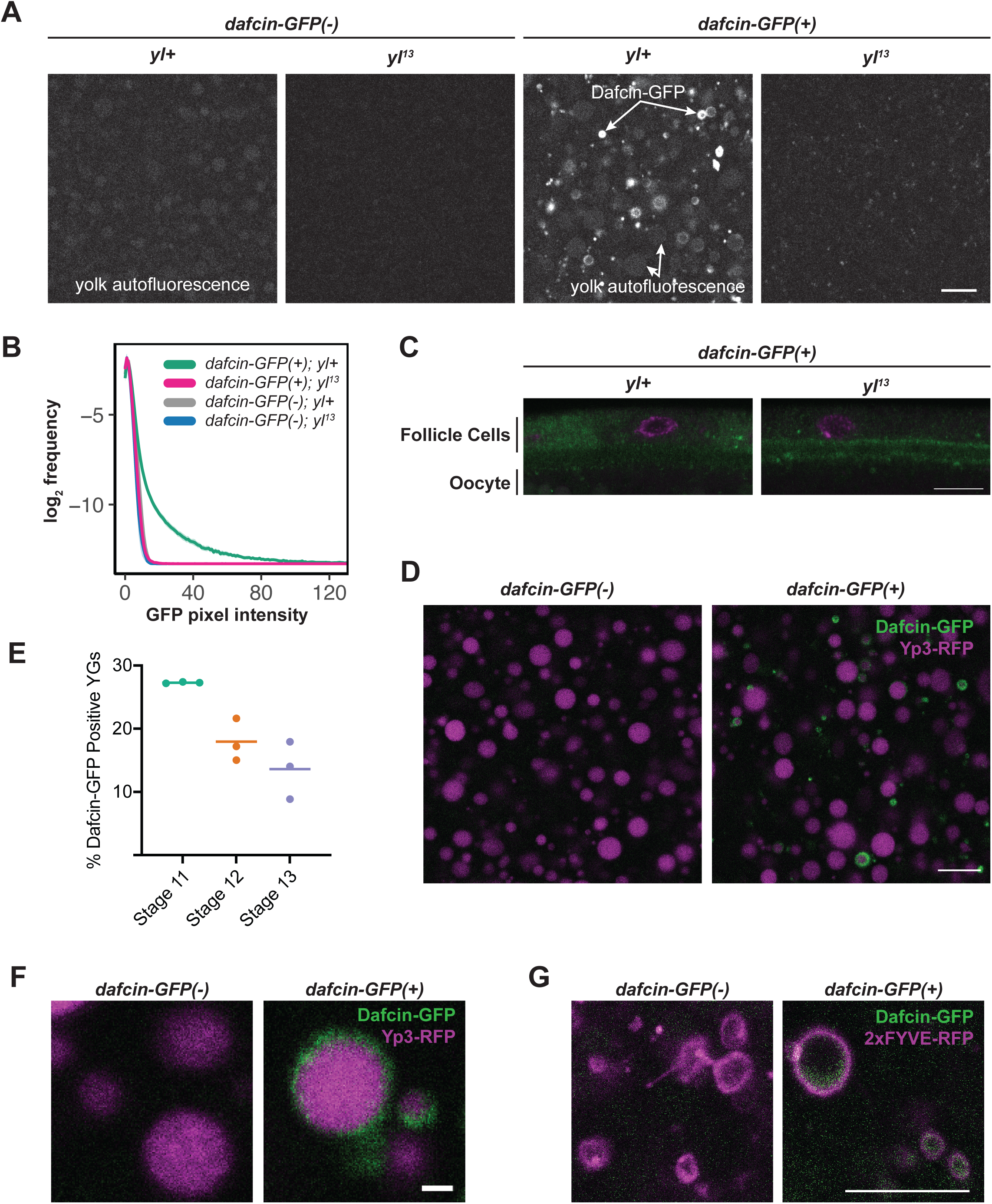
Dafcin-GFP enters the oocyte via endocytosis and localizes to a cortical layer within yolk granules. **(A)** Green fluorescence in stage 12 oocyte cytoplasm from females carrying either the endogenous *dafcin* gene (left panels) or *dafcin-GFP* gene (right panels). The females are either wildtype or mutant for the *yl* gene, as indicated. Autofluorescence from yolk granules is not detectable in *yl*^13^, consistent with a complete disruption of yolk granule biogenesis^46^. **(B)** Quantification of the analysis with *yl* mutants. Shown are frequency distributions of pixel-wise green fluorescence intensity in oocyte images. Moving line averages are in solid lines, and standard deviations are in shaded areas. Genotypes are as indicated. **(C)** Mid-sagittal section through *dafcin-GFP* stage 11 egg chambers, highlighting the follicle epithelium and oocyte cortex separated by a lumen. Egg chambers are either wildtype or mutant for the *yl* gene, as indicated. **(D)** Green fluorescence in stage 12 oocyte cytoplasm from females carrying either the endogenous *dafcin* gene (left panel) or *dafcin-GFP* gene (right panel). Shown also is Yp3-RFP localization (purple) to identify yolk granules. **(E)** Quantification of Yp3-RFP-positive yolk granules that are also positive for Dafcin-GFP. Each datapoint represents an independent oocyte sample at the indicated developmental stage. Bars represent the mean. **(F)** High magnification view of Yp3-RFP bodies (purple) and GFP fluorescence (green) in stage 12 oocytes from females carrying either the endogenous *dafcin* gene (left panel) or *dafcin-GFP* gene (right panel). **(G)** High magnification view of 2xFYVE-RFP-positive granules (purple) and GFP fluorescence (green) in stage 10B oocytes from females carrying either the endogenous *dafcin* gene (left panel) or *dafcin-GFP* gene (right panel). Scale bars represent 10 μm in (A), (C), (D), and (G), and scale bar represents 1 μm in (E).

Yolk proteins are synthesized in the fat body and follicle cells, and they are subsequently endocytosed by the oocyte where they are stored in specialized lysosomes known as yolk granules^46–48^. In contrast to many internalized ligands that are degraded in lysosomes, yolk proteins are stored in yolk granules for later use during embryogenesis. Yolk granules retain relatively high pH until embryogenesis, when the pH drops to levels more typical of lysosomes, and yolk proteins are degraded for nutrition^49–51^. Since Dafcin-GFP uptake by the oocyte is dependent on endocytosis of yolk proteins, we tested whether Dafcin-GFP is localized in yolk granules. There are three distinct yolk proteins (Yp1, Yp2, Yp3) found in yolk granules^52,53^. When yolk granules were visualized by an RFP-tagged form of Yp3, Dafcin-GFP was found to co-localize with Yp3-RFP in some of the yolk granules (Fig. 5D).

Oocytes accumulate yolk proteins over 22 hours, from the beginning of stage 8 to the end of stage 10B^54,55^. However, *dafcin* mRNA expression does not begin until stage 10B, which only lasts 4 hours but corresponds to a peak in endocytic activity. Therefore, we expected that only yolk granules formed during stage 10B would contain Dafcin-GFP, whereas yolk granules formed prior to that stage would lack it. As expected, only 27% of yolk granules contained Dafcin-GFP in stage 11 oocytes (Fig. 5E). This result is consistent with Dafcin-GFP specifically loading into stage 10B granules. Interestingly, the percent of Dafcin-GFP-positive yolk granules was further reduced at stages 12 and 13 (Fig 5E). One possibility is that the protein might be unstable in some maturing yolk granules.

Closer examination of yolk granules containing Yp3-RFP and Dafcin-GFP showed that the two proteins were segregated within the yolk granule. A large central body of Yp3 was surrounded by a cortical layer of Dafcin (Fig. 5F). This pattern may account for the rings of Dafcin-GFP in the oocyte cytoplasm seen earlier (Fig. 4A). We wondered if Dafcin-GFP was located inside or outside of the lipid membrane enveloping the yolk granule. To answer this question, we visualized the yolk granule membrane with 2xFYVE-tagged RFP, which inserts into membranes containing phosphatidylinositol 3-phosphate (PI(3)P), typically found in early endosomal membranes^56,57^. Each ring of Dafcin-GFP was localized inside a ring of 2xFYVE-RFP, indicating that Dafcin-GFP resides in a cortical layer interior to the yolk granule’s lipid membrane (Fig. 5G).

Electron microscopy had characterized yolk granules as having ultrastructural complexity^55,58^. Mature granules have a superficial layer of less dense material surrounding a central electron-dense body of yolk protein. The superficial layer lies beneath the yolk granule membrane. If comparable, it would suggest that Dafcin-GFP is located in this superficial layer.

### Dafcin regulates the size of yolk granules

Although Dafcin-GFP exhibits a striking mechanism of transport after its synthesis, it does not necessarily mimic the behavior of the endogenous microprotein. Numerous attempts to detect the microprotein by mass spectrometry were unsuccessful (data not shown). Thus, we could not be certain that the endogenous microprotein is transported to yolk granules. However, if Dafcin-GFP accurately mimics the transport properties of the microprotein, then we predict that Dafcin functions in some way to regulate yolk granules.

To understand the function of endogenous Dafcin, two independent frameshift mutant alleles were generated using CRISPR/Cas9 non-homologous end joining that targeted the start codon of the *dafcin* sORF (Fig. 6A). Both alleles are predicted to eliminate the start codon in *dafcin* mRNA isoforms 1 - 3 and block in-frame translation of the C-terminal 21 amino acids in *dafcin* mRNA isoform 4. Mutant flies were both viable and fertile under rich nutrient conditions.

**Figure 6.**
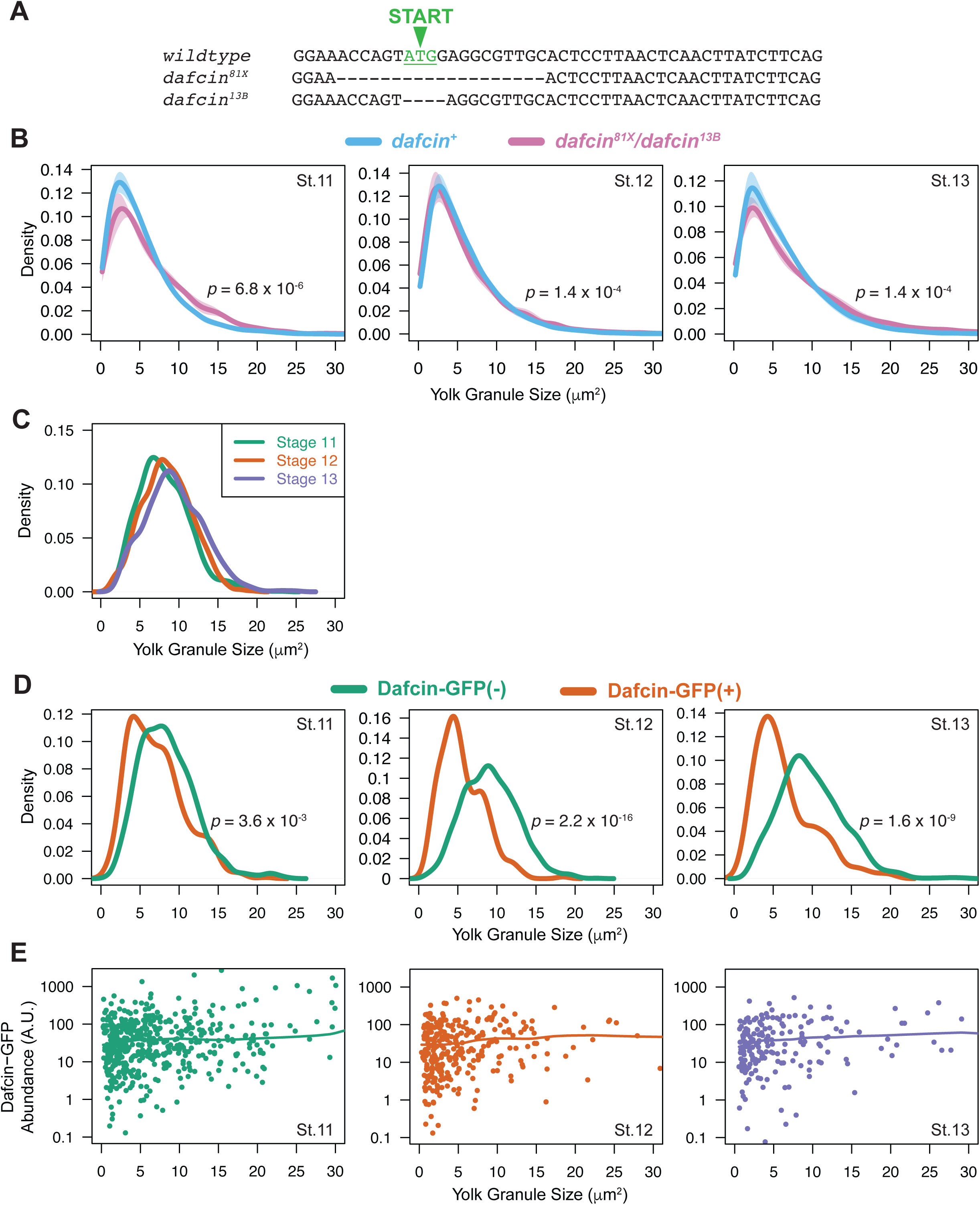
Dafcin limits the size of yolk granules in vivo. **(A)** Frameshift alleles *dafcin^81X^*and *dafcin^13B^* were generated using CRISPR/Cas9 non-homologous end joining. **(B)** Density distribution plots of Yp1-positive yolk granule size. Lines are moving line averages and shaded regions are standard deviations. Transheterozygous *dafcin* mutant females (*dafcin^81X^*/*dafcin^13B^*) have larger yolk granules than wildtype females at stages 11, 12, and 13. *P*-values shown are from Wilcoxon tests. **(C)** Density distribution plots of Yp3-positive yolk granule size. Shown are size distributions of yolk granules measured in oocytes of different stages. Kolmogorov-Smirnov tests return *p*-values above 0.10 comparing stage 11 to 12 and stage 12 to 13. However, tests return *p* = 0.003 comparing stage 11 to 13. **(D)** Density distribution plots of Yp3-positive yolk granule size in staged oocytes from wildtype females. Shown are size distributions of yolk granules classified as lacking Dafcin-GFP compared to those yolk granules containing Dafcin-GFP. *P*-values shown are from Kolmogorov-Smirnov tests. **(E)** Dafcin-GFP abundance in Yp3-positive yolk granules as a function of their size. These measurements are shown for staged oocytes from wildtype females. Each datapoint represents measurement of a single yolk granule. Lines represent moving line averages. Note the y-axis of each plot is in log_10_ scale.

Mutant oogenesis appeared normal, and yolk granules marked with Yp1-GFP were present in the oocytes (Supp. Fig. 3). However, the size of mutant Yp1-GFP yolk granules was significantly larger than those found in wildtype oocytes (Fig. 6B). The shift in the mutant yolk granule size distribution appeared to affect a subset of yolk granules. Although a difference in size was observed from stages 11 to 13, the size difference was greatest in stage 11 oocytes shortly after *dafcin* was first expressed (Fig. 6B). When we measured the size distribution of wildtype yolk granules as oocytes matured, we found a small but progressive increase in yolk granule size over developmental time (Fig. 6C). It would suggest that Dafcin slows down the maturation of yolk granules as they increase in size over time. This effect appears to limit the size of some but not all yolk granules.

We had observed Dafcin-GFP present only in 15 - 25% of yolk granules. If Dafcin limits the size of these yolk granules, then we expect that yolk granules containing Dafcin-GFP should be smaller than yolk granules lacking Dafcin-GFP. We compared the size distributions of Dafcin-GFP-positive and -negative yolk granules marked with Yp3-RFP (Fig. 6D). As expected, yolk granules containing Dafcin-GFP were smaller than those lacking Dafcin-GFP, and the difference became more pronounced as oocytes matured. This bias could also be seen when plotting Dafcin-GFP abundance within yolk granules (Fig. 6E). There were many more small yolk granules containing Dafcin-GFP than larger yolk granules, regardless of oocyte stage. Strikingly, the average abundance of Dafcin-GFP in yolk granules did not depend on yolk granule size. However, its abundance was highly variable between yolk granules, ranging over three orders of magnitude (Fig. 6E). Heterogeneity in Dafcin-GFP abundance was most pronounced in the smaller yolk granules.

### Dafcin microprotein limits liposome size in vitro

Dafcin is localized near the inner monolayer of yolk granule membranes, suggesting it may associate with the lipid membrane. As noted previously, Dafcin shares striking physico-chemical similarities with the influenza HA fusion peptide (Fig. 1F). When the virus enters a host cell via endocytosis, HA peptide inserts into the inner monolayer of the endosome membrane, initiating virus-host membrane fusion^29^. The peptide is able to insert due to the bulky hydrophobic surface on its amphipathic helix. Once inserted, HA peptide induces negative curvature of the membrane due to its acidic hydrophilic face^31^. We wondered if the common structural features of Dafcin and HA peptide might suggest that the two also have similar biochemical activities.

If HA peptide binds a lipid bilayer in vitro and induces negative curvature as the bilayer is forming a vesicle, then the peptide will stimulate the formation of smaller vesicles. This could be tested in vitro by forming liposomes in the presence of HA peptide. We generated multilamellar vesicle (MLV) liposomes composed of the unsaturated lipid 1,2-dipalmitoleoyl-*sn*-glycero-3-phosphoethanolamine (DPoPE). Pure HA peptide was incubated at a 1:200 molar ratio of peptide:lipid during liposome formation at pH 6.78, and the size of liposomes was measured. Compared to untreated liposomes, there was a significant reduction in liposome size when HA peptide was present (Fig. 7A, Supp. Fig. 4A). We repeated this experiment but incubated pure Dafcin microprotein with lipids at a 1:200 molar ratio. Strikingly, Dafcin stimulated a greater reduction in the size of liposomes than HA peptide (Fig. 7A, Supp. Fig. 4A). These effects were observed at a more acidic pH (pH 5.0) as well (Fig. 7B, Supp. Fig. 4B). This result strongly indicates that Dafcin interacts with lipid bilayers in a manner resembling HA peptide and provides a biochemical mechanism for Dafcin’s function to make smaller yolk granules in vivo (Fig. 7C).

**Figure 7.**
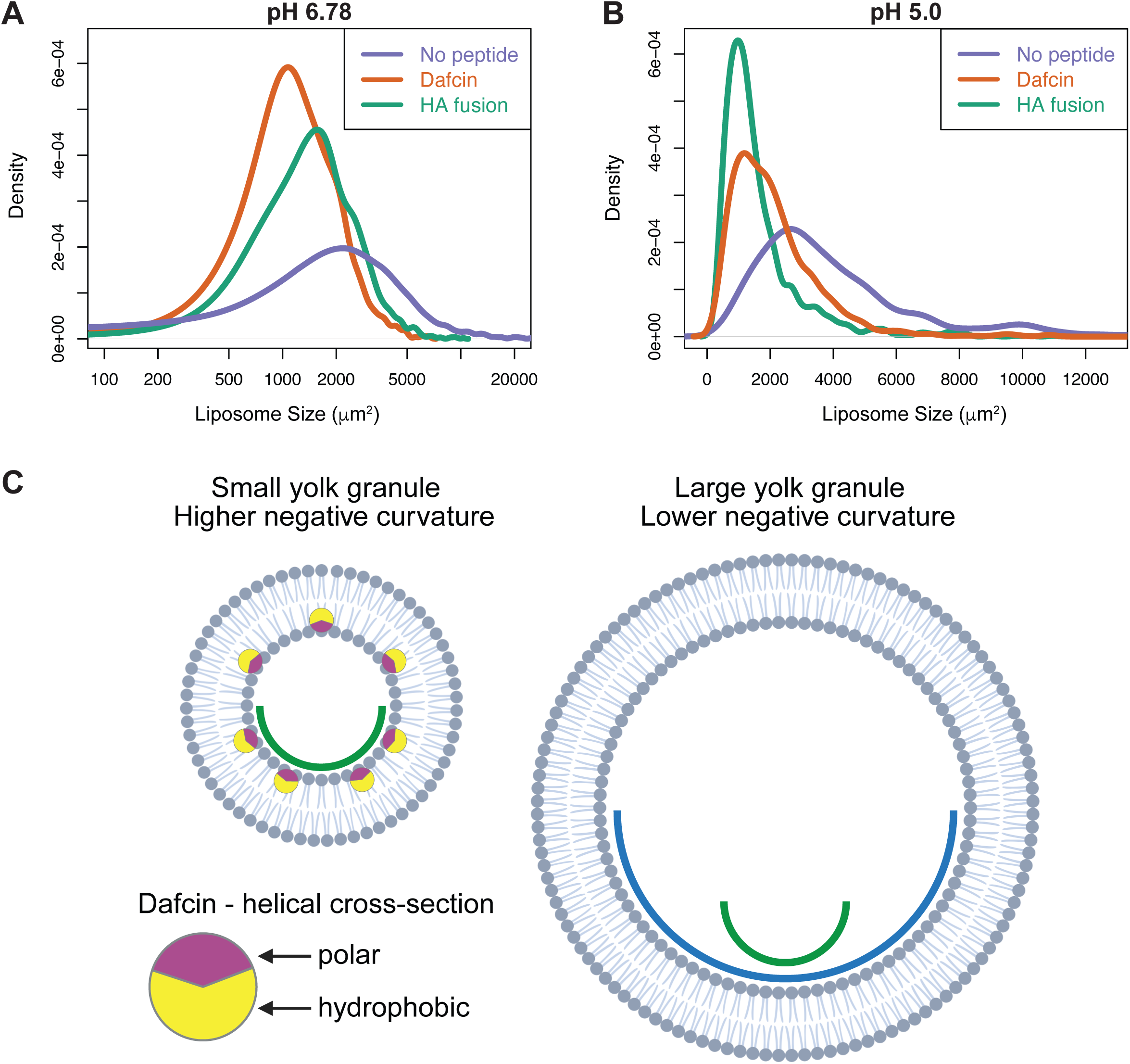
Dafcin interacts with lipid bilayers to restrict vesicle size. (A) Density distribution plots of DPoPE liposome size generated without peptide or in the presence of Dafcin or HA fusion peptide at pH 6.78. Kolmogorov-Smirnov tests return *p* < 2.2 x 10^-16^ and *p* = 1.3 x 10^-14^ when comparing control to Dafcin and HA fusion peptide, respectively. (B) Density distribution plots of DPoPE liposome size generated without peptide or in the presence of Dafcin or HA fusion peptide at pH 5.0. Kolmogorov-Smirnov tests return *p* < 2.2 x 10^-16^ when comparing control to both Dafcin and HA fusion peptide. (C) Working model for Dafcin’s interaction with higher negative curvature membranes found inside of small yolk granules. The hydrophobic face of Dafcin is depicted in yellow, with the polar face depicted in magenta. Created in BioRender. Nyberg, K. (2026) https://BioRender.com/fqd9bhy.

## Discussion

### The role of Dafcin in regulating yolk granule size

The striking physicochemical similarity between Dafcin and influenza HA fusion peptide at limiting lipid vesicle size suggests a compelling model for how Dafcin could regulate yolk granule size via its interactions with lipid bilayers. After uptake into the oocyte via endocytosis, the hydrophobic face of Dafcin’s alpha-helix inserts into the inner monolayer of the maturing yolk granule. Dafcin then stabilizes negative curvature of the yolk granule membrane, inhibiting the growth of maturing yolk granules.

What fitness benefit might Dafcin provide by ensuring some yolk granules are small? The vitellogenins or yolk proteins inside yolk granules are stored for later use as a nutrient source in the developing embryo. Catabolism of yolk proteins occurs as the yolk granules acidify, through the action of proton pumps, activating acid hydrolases that ultimately break down the complex yolk proteins into free amino acids^49,51^. The population of small yolk granules maintained by Dafcin have a higher surface area-to-volume ratio than larger yolk granules, owing to their spherical geometry. The increased surface-to-volume ratio might facilitate more rapid and complete acidification via the proton pumps and thus provide a more efficient nutrient source for embryogenesis. For females fed on nutrient-rich diets, the small yolk granules made by Dafcin may offer little benefit to the survival of their offspring since they would contain an abundance of yolk. For females fed on nutrient-poor diets, the small yolk granules made by Dafcin could provide their nutrients more efficiently and thus improve an embryo’s chance of survival.

Dafcin was initially identified as a potential microprotein through sequence conservation of its sORF across the *melanogaster* subgroup, indicating that it evolved under purifying selection. Even more interestingly, *dafcin* evolved ∼3.5-15 million years ago within the major DAFC on the X chromosome, which has also been shown to be amplified in the medfly *Ceratitis capitata*, suggesting that amplification at this locus is broadly conserved in *Drosophila*^59,60^. Perhaps *dafcin* is taking advantage of this unique genome environment in late-stage follicle cells to enhance or specify its expression.

### Trafficking potential of Dafcin

Dafcin is able to traffic a fused GFP cargo from the ovarian follicle cells into the yolk granules within the oocyte. It remains unclear how Dafcin exits the follicle cells. The 21-amino acid Dafcin microprotein lacks a canonical ER translocation signal sequence^32^. However, as shown by transgenic Dafcin-GFP expression, the 21-amino acid microprotein is sufficient for localization to the Golgi, exit from the follicle cells, and uptake into the oocyte. Thus, Dafcin may still enter the conventional secretory pathway at some point, though perhaps not through the ER.

Interestingly, transgenic Dafcin-GFP was not enriched near the apical domain of follicle cells, in contrast to apical enrichment observed with endogenous Dafcin-GFP. Since transgenic Dafcin-GFP was expressed at high levels within the follicle cells, this lack of enrichment could be a consequence of cytoplasmic saturation. Another possibility is that the 5’ UTR used in the *dafcin-GFP* transgene (isoform 2), which was abundant in RNA-Seq datasets but not our 5’ RACE clones, is not representative of the endogenous *dafcin* RNA population. The 5’ UTRs from isoforms 1 or 3 may direct protein translation in the apical cell domain while the 5’ UTR does not. It is also possible that the rare alternative isoform of Dafcin with an ER signal sequence (isoform 4) contributes in some degree to endogenous Dafcin-GFP localization.

While uptake of Dafcin-GFP into the oocyte largely occurred via receptor-mediated endocytosis dependent on Yl, it is not clear if Dafcin uptake is strictly dependent on Yl. Possibly, Dafcin can enter a cell via any type of endocytic event, but the predominant endocytic events in stage 10B oocytes are driven by Yl-mediated endocytosis^46^. In this situation, Dafcin would bind the outer monolayer of the cell membrane and be passively engulfed in invaginating endosomes driven by Yl binding to yolk protein. Several observations favor this latter explanation for Dafcin uptake. First, there is no evidence that *Drosophila* yolk proteins share any structural similarities with Dafcin, making it unlikely that the amphipathic Dafcin would function as a Yl ligand. Second, many amphipathic alpha-helices directly bind to lipid bilayers via their hydrophobic surfaces^15,16,40,41^. If Dafcin directly binds to lipid membranes, its endocytosis would be further stimulated if binding is stabilized by negative membrane curvature. Indeed, simulations of interactions between amphipathic alpha-helical peptides and lipid bilayers predict that amphipathic peptides with negative charges on their hydrophilic faces are more likely to interact with negatively curved membranes^61^. Third, in the absence of Yl-mediated endocytosis, Dafcin-GFP is trapped in the intercellular gap between the oocyte and follicle epithelium and is in close proximity to their cell membranes, suggesting that Dafcin has intrinsic affinity for interacting with lipid bilayers.

The localization of transgenic Dafcin-GFP to the Golgi of wing disc cells and S2 cells suggests that Dafcin’s remarkable trafficking ability is not restricted to follicle cells. The extent of Dafcin’s trafficking potential has just begun to be fully explored. If Dafcin’s secretion and uptake are driven largely through interactions with lipid bilayers, it is not unreasonable to presume that Dafcin could traffic cargoes between any number of eukaryotic cells, including mammalian cells. There is also no reason to believe that its cargo should be restricted to GFP. Indeed, Dafcin’s ability to traffic GFP between cells is not unlike that of cell penetrating peptides like HIV TAT and Penetratin, which have great potential as vehicles for drug delivery^62,63^.

### Figure Legends

**Supplementary Figure 1.**
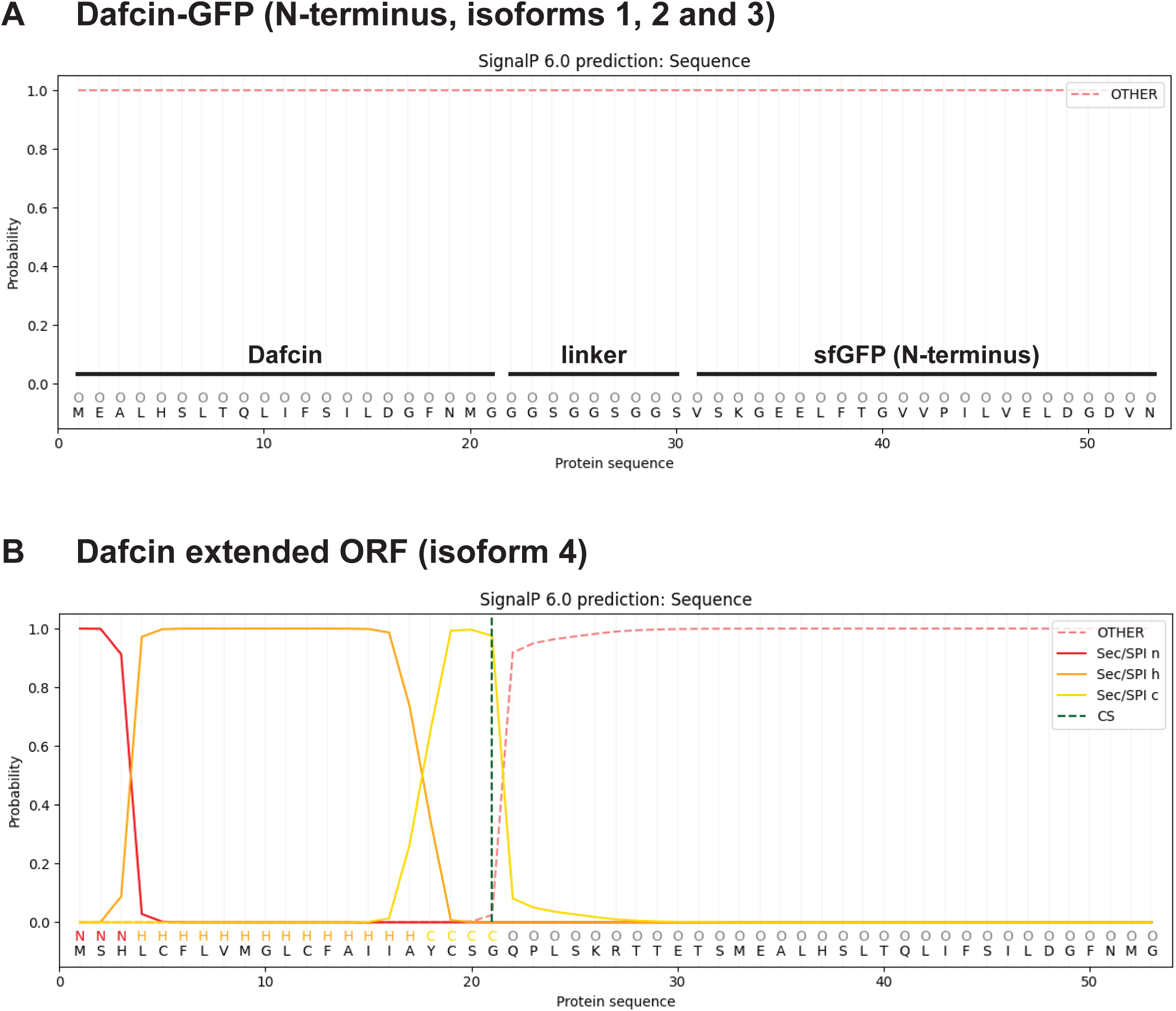
Signal peptide prediction of Dafcin using SignalP 6.0^67^. SignalP 6.0 signal peptide predictions of (A) the N-terminal 53 amino acids of Dafcin-GFP, including the entire Dafcin open reading frame, the flexible linker sequence, and the N-terminus of superfolder GFP, and (B) the extended open reading frame of *dafcin* isoform 4 (see Fig. 3C), with the likelihoods of each amino acid contributing to the n-region (“Sec/SPI n”), h-region (“Sec/SPI h”), or c-region (“Sec/SPI c”) of a signal peptide or a non-signal peptide region (“OTHER”) plotted on the y-axis. A predicted signal peptide cleavage site (“CS”) is indicated by a vertical dashed line. SignalP 6.0 was run using the following settings: organism = eukarya, output format = long, model mode = fast.

**Supplementary Figure 2.**
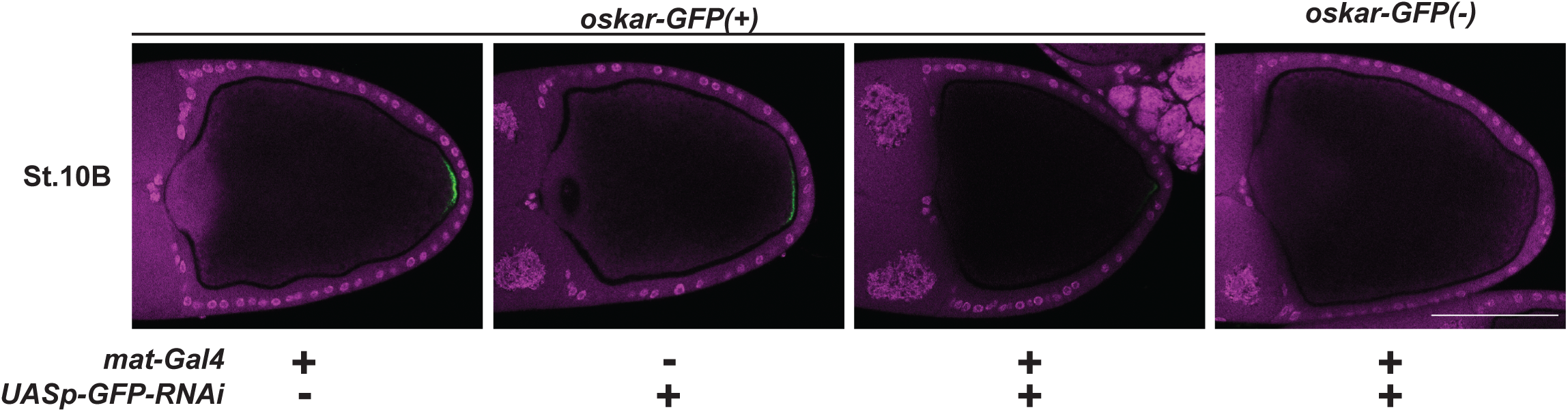
Validation of RNAi knockdown of germline *oskar-GFP*. RNAi knockdown of *oskar-GFP* expression using an inducible UAS-RNAi transgene driven by *maternal α-tubulin-Gal4* (*mat-Gal4*) in stage 10B oocytes. Scale bars represent 100 μm.

**Supplementary Figure 3.**
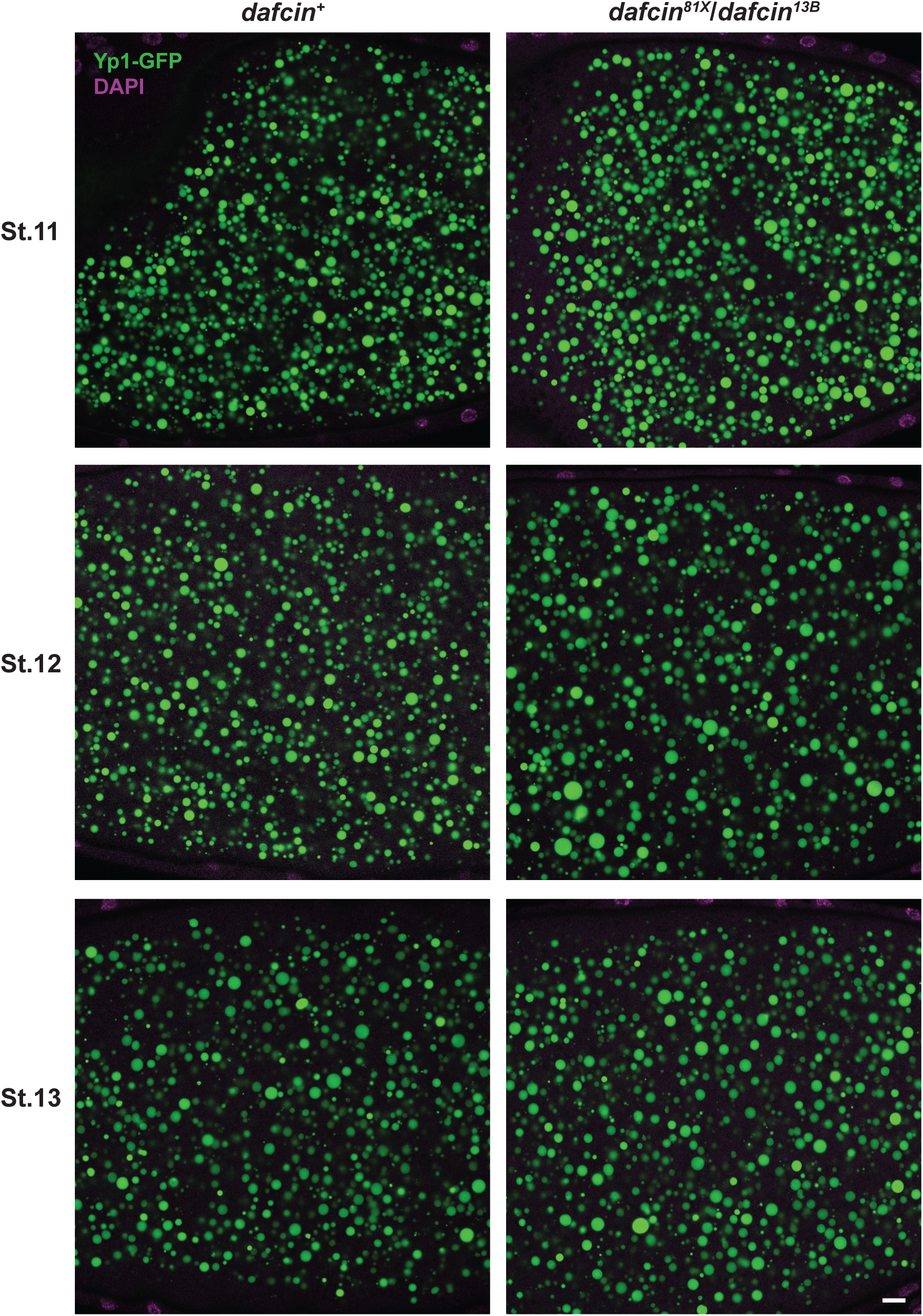
Yolk granules are present in both *dafcin* wildtype and mutant egg chambers. Yolk granules marked with Yp1-GFP in Stage 11, Stage 12, and Stage 13 *dafcin^+^* and *dafcin^81X^*/*dafcin^13B^*egg chambers. Scale bar represents 10 μm.

**Supplementary Figure 4.**
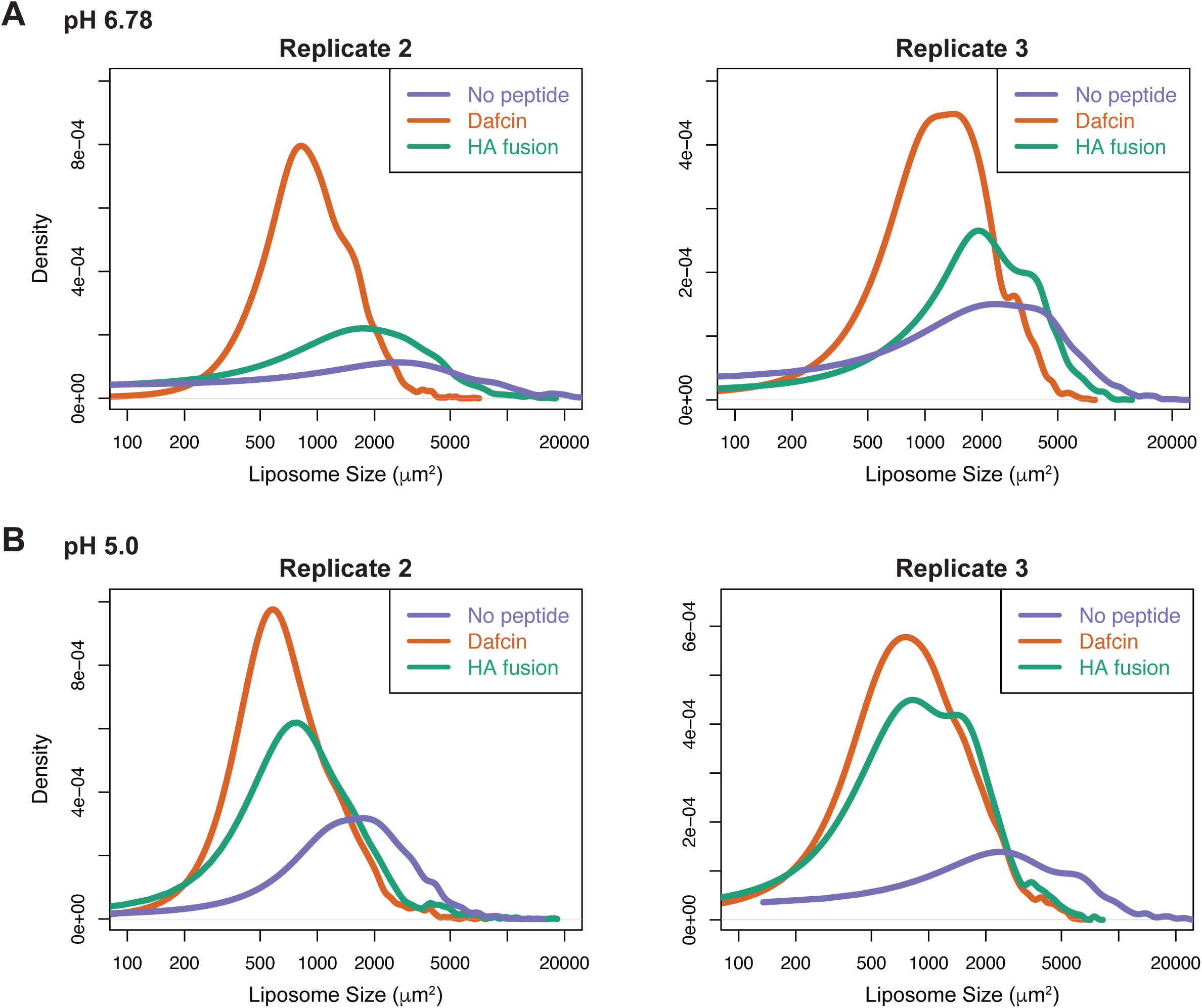
Dafcin interacts with lipid bilayers to restrict vesicle size. Additional replicates of in vitro experiments depicted in Figs. 7A and 7B. (A) Density distribution plots of DPoPE liposome size generated without peptide or in the presence of Dafcin or HA fusion peptide at pH 6.78. Kolmogorov-Smirnov tests return *p* < 2.2 x 10^-16^ when comparing control to Dafcin in both replicates and *p* < 2.2 x 10^-7^ and *p* = 0.00182 when comparing control to HA fusion peptide in replicates 2 and 3, respectively. (B) Density distribution plots of DPoPE liposome size generated without peptide or in the presence of Dafcin or HA fusion peptide at pH 5.0. Kolmogorov-Smirnov tests return *p* < 2.2 x 10^-16^ when comparing controls to both Dafcin and HA fusion peptide in both replicates.

**Supplementary File 5. RNA smFISH probe sets for *dafcin* and *CR44289*.**

**Supplementary File 6. Ovary RNA-Seq datasets from multiple *Drosophila* species used in this study.**

**Supplementary File 7. Sequence file of *dafcin-GFP* donor repair plasmid for CRISPR/Cas9 homology-directed repair in GenBank format.**

**Supplementary File 8. Sequence file of *UAST-dafcin-GFP* plasmid for PhiC31 transgenesis in GenBank format.**

## Methods

### Characterization of *dafcin* RNA in *D. melanogaster* and other species of *Drosophila*

The *dafcin* gene (FlyBase annotation symbol: *CR43834*, FlyBase gene ID: FBgn0264382) is annotated as a long non-coding RNA that produces a single RNA transcript (FlyBase transcript ID: FBtr0332277) in the most recent *Drosophila melanogaster* genome assembly (Berkeley *Drosophila* Genome Project r6.67). *Dafcin* RNA is enriched in the ovaries in multiple tissue-specific RNA-Seq datasets (modENCODE and FlyAtlas2, both accessed via FlyBase)^68,69^.

To capture full-length *dafcin* RNA transcripts, both 5’ and 3’ RACE (rapid amplification of cDNA ends) were performed on RNA extracted from *D. melanogaster* ovaries. All flies were reared on standard molasses cornmeal media. To stimulate embryo production, *w*^1118^ females were crossed to males of the same line and fed with yeast paste. After 4-6 days, ovaries were isolated from females fed with yeast paste and placed directly into TRIzol reagent. Total RNA was then extracted using the manufacturer’s protocol. 5’ RACE was then performed using the Invitrogen GeneRacer Kit. 3’ RACE was performed using a previously published protocol^70^. RACE products from each reaction were cloned using the Zero Blunt TOPO PCR Cloning Kit and sequenced.

Single-molecule RNA fluorescence in situ hybridization (RNA smFISH) was used to detect *dafcin* and the lncRNA *CR44289* in *D. melanogaster w*^1118^ ovaries using a previously described approach^71–74^. The probe set for *dafcin* RNA consisted of 27 non-overlapping oligonucleotides that are antisense to the mature *dafcin* transcript sequence. The *dafcin* probe set was designed manually using OligoAnalyzer (Integrated DNA Technologies) with predicted melting temperatures for each oligonucleotide between 68 and 71°C in 660 mM NaCl. The probe set for *CR44289* consisted of 48 non-overlapping oligonucleotides that are antisense to the mature *CR44289* transcript sequence designed using the Stellaris Probe Designer (Biosearch Technologies). RNA smFISH probe sets are listed in Supplementary File 5. Prepped samples were imaged on a Leica TCS SP8 confocal microscope.

To determine whether *dafcin* RNA expression was conserved in multiple species of *Drosophila*, *dafcin* RNA levels were estimated from ovary RNA-Seq datasets, including datasets from *D. melanogaster*, *D. simulans*, *D. yakuba*, *D. ananassae*, *D. pseudoobscura*, *D. willistoni*, *D. virilis*, *D. mojavensis*, and *D. grimshawi*. All analyzed datasets are listed in Supplementary File 6 and were obtained from the NCBI Sequence Read Archive using fastq-dump (SRA Toolkit v2.8.1)^75,76^, with the exception of two *D. pseudoobscura* datasets, DRR055260 and DRR055261, which were obtained from the National Institute of Genetics DNA Data Bank of Japan^77^. Read quality was assessed using FastQC v0.11.5^78^. Reads with flagged quality issues were filtered for base quality using IlluQC_PRLL.pl from the NGS QC Toolkit (v2.3.3)^79^ with default parameters and trimmed using TrimGalore v0.4.3^80^. High-quality reads were aligned to species-specific genome assemblies listed in Supplementary File 6 using STAR v2.6.0^81^ with the following parameters: --twopassMode Basic --outFilterMismatchNmax 2 --outFilterMismatchNmax 999 --outFilterMismatchNoverReadLmax 0.03 --outFilterScoreMinOverLread 0.3 --outFilterMatchNminOverLread 0.3. For *D. melanogaster* datasets, read counts for each gene from the dmel-all-r6.19.gtf annotation file were calculated using htseq-count v0.7.2 with union mode^82^. To estimate read counts for datasets from all other *Drosophila* species, the coordinates from the dmel-all-r6.19.gtf file were first converted from FlyBase r6 to FlyBase r5 using the liftOver tool^83^ from the UCSC genome browser with the dm6ToDm3.over.chain chain file and - minMatch=0.9. This converted r6.19 annotation file was then converted to coordinates of all other *Drosophila* species using the appropriate chain files listed in Supplementary File 6 and - minMatch=0.1. Read counts were then calculated from each species using the appropriate converted r6.19 annotation files. Read counts were then normalized to counts per million (cpm). The *dafcin* gene region was not alignable in species outside the *melanogaster* species subgroup.

### Predictions of *dafcin’s* coding potential

The coding potential of *dafcin* was predicted using PhyloCSF^39^. A multiple sequence alignment of the *dafcin* transcript FBtr0332277 in maf format was extracted from the *Drosophila* 27 species multiple genome alignment obtained from the UCSC genome browser^84^. Non-*Drosophila* species were removed from this alignment, and the maf alignment was then converted to a multiple fasta alignment using the AlignIO module in BioPython v1.85^85^. PhyloCSF was then run using the *dafcin* multiple fasta alignment as input along with the following options: 20flies --minCodons=10 --frames=3 --allScores --orf=ATGStop --removeRefGaps. The PhyloCSF score of 51.9710 decibans indicated that the *dafcin* ORF is ∼100,000-fold more likely to be protein-coding than non-coding.

### Structural predictions of the Dafcin microprotein

The alpha-helical structure of the Dafcin microprotein (MEALHSLTQLIFSILDGFNMG) was predicted using both AlphaFold 3^65^ and QUARK^86^, both available online. Amino acids were colored according to biochemical characteristics using PyMOL v3.1.6.1^64^, with hydrophobic residues depicted in yellow and polar (i.e., hydrophilic) residues depicted in magenta. Physicochemical properties of the Dafcin and HA fusion peptide (GLFGAIAGFIEGGWTGMIDGWYG) alpha-helices were predicted using HeliQuest v2, available online^31,66^. Dafcin’s potential as a signal peptide was predicted using the online version of SignalP 6.0 (“Eukarya” signal peptide models, “Fast” model mode)^67^.

### Generation of *dafcin-GFP* via CRISPR/Cas9 HDR

Flexible linker and superfolder GFP (GFP) sequences were inserted at the 3’ end of the endogenous *dafcin* 63-nucleotide ORF on the X chromosome in *w*^1118^ flies using a previously described CRISPR/Cas9 protocol^87,88^. A single guide RNA (sgRNA) was designed to target a PAM site at the 3’ end of the *dafcin* ORF (5’- CTTATCTTCAGCAT<u>TC</u>TGGATGG-3’), and the cleavage efficiency of this sgRNA was verified using a T7EI assay as previously described. A donor repair plasmid was constructed via Gibson assembly (HiFi DNA Assembly Master Mix, New England Biolabs) using *Eco*RV-linearized pBS-*GMR-eya*(shRNA) as the backbone. This donor plasmid also contained the *dafcin-linker-GFP* sequence, a removable scarless DsRed marker gene, and ∼1000 bp homology arms (Supplementary File 7). The two underlined nucleotides in the sgRNA target sequence were synonymously mutated from TC to CT in the donor plasmid to prevent targeting and cleavage of the donor repair plasmid by the Cas9/sgRNA RNPs: CTTATCTTCAGCAT<u>CT</u>TGGATGG. Injection of Cas9/sgRNA RNPs and screening of genetically modified flies were then performed as previously described^87,88^.

### Generation of transgenic *dafcin-GFP* via PhiC31 transformation

Transgenic flies containing *dafcin-GFP* under the control of the UAS enhancer were generated via PhiC31 transformation^89,90^. Briefly, the *dafcin-GFP* sequence, containing the *dafcin* mRNA isoform 2 (5’ UTR, 3’ UTR, and sORF), as well as the linker and superfolder GFP sequences, was inserted into *Eco*RI-linearized pUAST-attB (*Drosophila* Genomics Resource Center #1419) using Gibson assembly (Supplementary File 8)^91,92^. This plasmid was then injected into embryos containing both *nanos-PhiC31* integrase (BDSC #34770)^90^ and an attP site at the VK22 locus on the 2^nd^ chromosome (BDSC #9740)^93^. Surviving injected flies were crossed with the *w*^1118^ line, and the resulting G_1_ progeny were screened for presence of the mini-*white* gene, indicating successful integration of the transgene into the VK22 site. Transformed G_1_ individuals were then crossed to *CyO,dfd-YFP* balancer flies to generate a balanced stock^94^. To express the transgenic Dafcin-GFP in the ovarian follicle cells, flies carrying the *UAST-dafcin-GFP* transgene were crossed to flies carrying the *c825-Gal4* driver (BDSC #6987), which expresses Gal4 in late-stage follicle cells but not the female germline^95^.

### Confocal microscopy of Dafcin-GFP

To visualize both endogenous and transgenic Dafcin-GFP in fixed ovaries, newly eclosed females were cultured with males and fed with yeast paste to stimulate oogenesis. After 3-5 days, ovaries were dissected in cold PBS, individual ovarioles were gently teased apart using tungsten wires, and ovaries were fixed in 4% paraformaldehyde for 20 - 30 min with gentle rocking at room temperature. Ovaries were then washed once with PBS supplemented with 0.3% Triton X-100 (PBTx) for 5 min, stained with DAPI (2.5 μg/mL in PBTx) for 5 min at room temperature to visualize nuclei, and then washed three more times in PBTx for 5 min each, all with gentle rocking. Ovaries were then mounted in 30-40 μL Vectashield Plus (Vector Laboratories #H-1900) with a No. 1.5 coverslip (Zeiss) and imaged on a Leica TCS SP8 confocal microscope.

To visualize transgenic Dafcin-GFP in fixed wing discs, wandering 3^rd^-instar larvae carrying both the *UAST-dafcin-GFP* transgene and the *dpp.blk1-Gal4* driver (BDSC #93385)^96,97^ were dissected. Wing discs were fixed for 20 minutes in 4% paraformaldehyde solution and washed three times with PBTx and once with DAPI (2.5 μg/mL in PBTx). They were mounted in 40 μL of Vectashield Plus between two coverslips.

To detect co-localization of Dafcin-GFP with Golgi proteins, Dafcin-GFP (either endogenous or transgenic) tissues were isolated and fixed as described above. Fixed tissues were washed three times in PBTx for 5 min each at room temperature with gentle rocking. Tissues were blocked in 5% normal animal serum (NAS) in PBTx for 60-90 min at room temperature before overnight incubation with primary antibodies in PBTx + 5% NAS at 4°C with gentle rocking. Tissues were washed three times in PBTx for 5 min each at room temperature with gentle rocking and incubated for 1-2 hours with secondary antibody in PBTx + 5% NAS at room temperature with gentle rocking. Tissues were washed, stained with DAPI (2.5 μg/mL in PBTx), and mounted in Vectashield Plus as described above. To detect GMAP and Golgin245, tissues were blocked in 5% normal donkey serum, incubated in the primary antibody (1:2000, both raised in goat and available from DSHB)^98^, and then incubated in donkey anti-goat Alexa Fluor 633 secondary antibody (1:500, ThermoFisher #A-21082). To detect GM130, tissues were blocked in 5% normal goat serum, incubated with rabbit anti-GM130 (1:500, Abcam #Ab30637), and then incubated in goat anti-rabbit Alexa Fluor Plus 647 secondary antibody (1:500, ThermoFisher #A-32733). Imaging was performed on a Leica TCS SP8 confocal microscope.

To detect co-localization of Dafcin-GFP with yolk granules, yolk granules were marked by driving a *UAST-Yp3-mRFP* transgene (a gift from Stefan Luschnig)^99^ with the *r4-Gal4* driver (BDSC #33832), which expresses Gal4 in the fat body. To detect co-localization of Dafcin-GFP with early yolk granule membranes, membranes were marked by driving a *UASp-TagRFP-2xFYVE* transgene (a gift from Graydon Gonsalvez)^56^ with the *maternal α-tubulin-Gal4* driver (BDSC #7063), which expresses Gal4 in the female germline. Ovaries were then prepared as described above.

### Detection of Dafcin-GFP in *Drosophila* S2 cells

To test the translation potential of *dafcin,* a *Drosophila* S2 cell expression plasmid was constructed to express a Dafcin-GFP fusion protein, with EGFP fused to Dafcin’s C-terminus. To construct this, the stop codon within the ORF in *dafcin* isoform 2 was removed, *EGFP* sequence was inserted at the 3’ end of the *dafcin* ORF, and this fusion protein sequence, including the isoform’s 5’ and 3’ UTRs, was inserted into the *pMK33* expression plasmid^100^ via Gibson assembly. A control plasmid that retained the native *dafcin* stop codon was constructed in parallel, as was a transfection control plasmid that expressed *DsRed*.

Plasmids were transiently transfected into *Drosophila* S2* cells as follows. First, 5 mL of S2* cells in Schneider’s media (Gibco #21720024) supplemented with 10% fetal bovine serum (Gibco #10082147) and 1% penicillin/streptomycin (Gibco #15140122) were cultured in flasks at a density of 2 x 10^6^ cells/mL and incubated overnight. On the second day, 500 ng of each plasmid was transfected into cells using Effectene Transfection Reagent (Qiagen #301425) using standard protocol. On day three, expression of genes under the control of the metallothionein promoter was induced by adding CuSO_4_ to a final concentration of 0.5 mM. On day four, 1 mL of cells from each treatment was incubated on coverslips covered with 0.1% w/v poly-L-lysine for 2 hours, washed twice with cold PBS, and fixed in 4% paraformaldehyde in PBS for 10 min. Fixed cells were then washed and mounted in Vectashield Plus and imaged on a Zeiss Axioplan microscope.

### RNAi knockdown of Dafcin-GFP

RNAi knockdown of endogenous Dafcin-GFP was used to verify that oocyte-localized Dafcin-GFP originated in the follicle cells. Knockdown of Dafcin-GFP in the follicle cells was achieved using a *UAST-GFP-RNAi* transgene (BDSC #41553) driven by the *c825-Gal4* driver. Knockdown of Dafcin-GFP in the female germline was achieved using a *UASp-GFP-RNAi* transgene (BDSC #41551) driven by the *maternal _α_-tubulin-Gal4* driver. Both *GFP-RNAi* transgenes target the same sequence of GFP. RNAi knockdown of GFP in the female germline with the *UASp-GFP-RNAi* transgene was verified by targeting *oskar-GFP* (VDRC #318897) for knockdown (Supplementary Figure 2)^101^.

Ovaries from RNAi knockdown experiments were dissected, prepared, and imaged as described above. Pixel intensities of GFP signal for all pixels within regions of interest (ROIs) in follicle cells or oocytes were obtained using the histogram function in FIJI with 256 bins^102^. The frequency of each pixel intensity within the ROI was then calculated, log_2_-transformed, and plotted using the ggplot2 package in R^103^. Statistically significant differences between these distributions were assessed using the Kolmogorov-Smirnov test with Benjamini-Hochberg adjustment for multiple comparisons.

### *yolkless* mutant experiments

To determine whether the oocyte internalization of Dafcin-GFP was dependent on Yolkless-mediated endocytosis, endogenous *dafcin-GFP* flies were crossed to flies carrying a strong *yolkless* allele (*yl*^13^, BDSC #4320)^46,104^. Since both genes are on the X chromosome, dafcin-GFP was recombined with *yl*^13^ on the X chromosome. Potential recombinants were balanced over an *FM7c* balancer chromosome (BDSC #7756) and were screened for the presence of *dafcin-GFP* via PCR. The presence of *yl*^13^ was then tested by complementation crosses with the *yl*^17^ allele (BDSC #59958)^46^. Ovaries from *dafcin-GFP*, *yl*^13^ females were then prepared and imaged as described above and compared to ovaries from the parental stocks as controls.

### *dafcin* mutant generation and analysis

CRISPR/Cas9 mediated non-homologous end joining (NHEJ) was used to create multiple *dafcin* frameshift alleles using a sgRNA targeting the *dafcin* start codon. A sgRNA targeting a sequence near the start codon (5’-GACAACGGAAACCAGTATGG-3’) was inserted into the *pU6-BbsI-chiRNA* plasmid (DGRC #1362), which drives expression of the sgRNA using a Pol III promoter and terminator^105^. The sgRNA plasmid DNA was then injected into *vasa-Cas9* (BDSC #51324) embryos. Approximately 300 embryos were injected, and the 120 surviving G0 flies were crossed to *FM7c* to generate balanced stocks. Genomic DNA was extracted from balanced G_1_ lines, and potential frameshift mutations were detected via PCR of the *dafcin* gene region (Forward: GGGTATCACAGAAAGTCCTTA, Reverse: ATGCACTAATTTGATGTAATATGC) followed by Sanger sequencing. Transheterozygotes of the *dafcin^81X^* and *dafcin^13B^* frameshift alleles were used for all mutant analyses, while balanced G_1_ lines without frameshift mutations were used as wildtype controls.

To determine the effect of *dafcin* on yolk granule size, yolk granules in *dafcin^81X^*/*dafcin^13B^* transheterozygotes and wildtype controls were marked using a Yp1-GFP fusion protein (Vienna *Drosophila* Resource Center #318746)^101^. Three biological replicates of each genotype were imaged and analyzed. Ovaries were dissected from 3-day-old mated females in Schneider’s medium supplemented with 15% fetal bovine serum and fixed in 4% paraformaldehyde in PBS for 20 min at room temp with gentle rocking. Fixed ovaries were then washed, stained with DAPI, and mounted in Vectashield Plus as previously described. Yp1-GFP+ yolk granules in *dafcin^81X^*/*dafcin^13B^*and wildtype egg chambers were imaged on a Leica TCS SP8 confocal microscope. Single 2048 x 2048 pixel images were captured from stage 11, 12, and 13 egg chambers at uniform depth beneath the follicle cell layer (15 μm for stage 11; 20 μm for stages 12 and 13).

Initial pixel classification segmentation of yolk granules in these images was performed using Ilastik v1.4.0^106^. Segmentation was trained on three images from each genotype (six total). All potential features were considered, and yolk granule and background pixels were manually designated in the training set. Simple segmentation predictions were then exported from Ilastik as hdf5 files and imported into FIJI^102^. Within FIJI, pixel intensities were set to a minimum of 1 and a maximum of 2 under the Brightness/Contrast settings. A default black and white threshold was then applied under the Threshold settings. A watershed algorithm was then used to separate overlapping yolk granules. To reduce the number of segmented artifacts, segmented objects were then filtered for size and circularity, retaining objects between 25 and infinity pixels and with circularity between 0.7 and 1.0. Between 569 and 1,722 yolk granules were segmented within each egg chamber, and sizes of these segmented yolk granules were then measured. Distributions of yolk granule size were plotted using ggplot2 and analyzed using the Kolmogorov-Smirnov test in R^103^.

### Yolk Granule Quantification and Analysis

#### Segmentation

For each oocyte, optical slices were chosen for imaging and analysis in FIJI^102^. Yp3-RFP images were smoothed by a Gaussian blur of two pixels to prevent spurious segmentation. Next, Yp3-RFP objects were identified by thresholding based on the RFP channel intensity, and a Watershed algorithm was applied. Segmentation parameters were optimized to obtain objects with at least 20 pixels. Errors remaining after segmentation were manually corrected, and area and centroid data were collected.

Next, for each RFP object so identified, they were dilated by four pixels in order to reliably capture Dafcin-GFP signal, which typically resides in a thin cortex around the Yp3-RFP signal. We previously determined that virtually all Dafcin-GFP signals are located within four pixels of a Yp3-RFP boundary. Once dilation was complete, the Watershed algorithm was run again, and any errors were manually corrected. Segmentation parameters were reapplied to capture objects of at least 300 pixels. The Dafcin-GFP image was then analyzed to measure green fluorescence intensities in the dilated objects, whose areas and centroids were also measured. Data from nondilated red objects and dilated green objects were automatically matched based on the XY coordinates of the objects’ centroids. Matching errors were manually corrected to generate a robust and accurate dataframe.

#### Autofluorescence correction

A majority of imaged yolk granules did not contain Dafcin-GFP signal and therefore displayed background levels of green autofluorescence due to yolk protein. Dafcin-GFP-positive yolk granules were present in the right-hand tail of the green fluorescence distribution. Therefore, we imaged in parallel, under identical conditions, stage-matched oocytes from animals expressing Yp3-RFP but lacking the *dafcin-GFP* transgene. Images were processed as described above to obtain a population of green fluorescence intensity values taken from the dilated objects.

We fit a Gaussian distribution to the population of intensity values, using maximum-likelihood estimation, and finding the mean and standard deviation of that fit. In order to separate Dafcin-GFP positive yolk granules, we chose a cut-off percentile based on the normal distribution of autofluorescence, below which yolk granules were deemed to be Dafcin-GFP negative. We set this cut-off at the 95th percentile for all analyses. Only objects with values above the threshold were assumed to be Dafcin-GFP positive. Thus, to normalize measurements across all stage-matched oocyte samples, this cut-off value was subtracted from the measured fluorescence intensities for all segmented objects in that sample.

#### Dafcin-GFP fluorescence scaling

To measure the total abundance of Dafcin-GFP in a yolk granule, the mean green fluorescence intensity of a Dafcin-GFP-positive object was multiplied by the area (in pixels) of the dilated object. This value was then normalized to μm^2^ by dividing total fluorescence by 123 (the number of pixels in 1.0 μm^2^).

#### Sampling

For each stage of oogenesis analyzed, three replicate oocytes were sampled. The number of yolk granules segmented in each oocyte were similar to one another. The total number of yolk granules analyzed were: 1,835 for stage 11; 1,699 for stage 12; 1,431 for stage 13.

### Preparation of Liposomes

Liposomes were prepared according to previously published methods^31,107,108^. DPoPE (1,2-dipalmitoleoyl-sn-glycero-3-phosphoethanolamine, Avanti #850706) was dissolved at 5.5 mg/mL in a 2:1 (v/v) chloroform:methanol mixture. Lipid solutions were handled exclusively in borosilicate glass vials and stored in glass vials fitted with PTFE-lined caps. Organic solvents and solutions were transferred using glass pipettes. The lipid stock solution was overlaid with nitrogen gas, tightly capped, and stored at −20 °C in an evacuated desiccation chamber containing Drierite.

Dafcin and HAfp23 peptides were synthesized by AnaSpec Inc. (≥90% purity), supplied as lyophilized powders, and used without further purification. Peptides were weighed to achieve an approximate final peptide:lipid molar ratio of 1:200 and transferred into a small borosilicate glass vial. An aliquot (400 μL) of the 5.5 mg/mL DPoPE solution was added to the peptide-containing vial, and the peptide was dissolved by gentle pipetting.

The lipid–peptide mixture was dried to a thin film under a gentle stream of nitrogen gas for 3 hours. Once visibly dry, vials were capped, flash-frozen in liquid nitrogen, immediately loosened, and transferred to a lyophilizer flask. Samples were subjected to overnight lyophilization to remove residual organic solvents. During lyophilization, vials were wrapped in aluminum foil to protect samples from light exposure.

Acetate buffer (10 mM sodium acetate, 150 mM NaCl, 1 mM EDTA) was prepared in Milli-Q water and adjusted to pH 5.0 or 6.78 using 0.1 M acetic acid or NaOH. Prior to hydration, both the dried lipid films and acetate buffer were equilibrated at 37 °C for 10 minutes. Pre-warmed buffer (200 μL) was added directly to the lipid film. Samples were vortexed briefly, sealed with Parafilm, and incubated at 37 °C for 30 minutes to facilitate hydration.

Hydration was enhanced using repeated vortex–incubation cycles consisting of 30 s vortexing followed by 30 s incubation at 37 °C; this cycle was repeated nine times (total vortex time: 5 min). The hydrated lipid suspension was then subjected to freeze–thaw cycling to promote vesicle homogeneity. Samples were flash-frozen in liquid nitrogen and rapidly thawed in a 37 °C water bath, followed by brief vortexing. This freeze–thaw cycle was repeated six times.

### Liposome Quantification and Analysis

#### Imaging

Immediately after liposomes were prepared, aliquots of the liposome suspension were mounted for dark-field microscopy using a Zeiss Axioplan microscope. Images were captured as CZI files with a Zeiss Axiocam ERc 5s using ZEN Blue 3.1 imaging software. Each image had dimensions of 2560 x 1920 pixels, and each pixel intensity was recorded in 8-bit RGB mode. CZI files were converted to TIFF format using the ZEN Blue software and converted to grayscale.

#### Segmentation and measurement

For each TIFF file, liposomes were identified by thresholding based on pixel intensity. Segmentation parameters were optimized to obtain masked objects. Errors remaining after segmentation were manually corrected. The areas of all corrected and segmented objects were then recorded.

Three replicate experiments were performed for liposome formation at pH 6.78 and another three replicate experiments for liposome formation at pH 5.0. In each experiment, liposomes were formed in the presence of HA fusion peptide, in the presence of Dafcin peptide, and in the absence of either peptide. For each treatment, a minimum of 10 microscope images were captured per experiment. This enabled us to measure between 608 and 2,704 liposomes for each replicate and treatment.

## Acknowledgments

We thank Jessica Hornick and the Biological Imaging Facility at Northwestern (RRIS:SCR_017767) for confocal microscopy and the Keck Biophysics Facility, a shared resource of the Robert H. Lurie Comprehensive Cancer Center of Northwestern University supported in part by the NCI Cancer Center Support Grant #P30 CA060553, for instrumentation. We also thank Hiroaki Sai, Leonel Bustamante-Carballo, and Justin Lorieau for guidance working with HA fusion peptide and liposomes.

## Funding

This work was supported by the National Institutes of Health (R35GM118144 to RWC and F32GM122349 to KGN) and undergraduate research grants to RE and CWPS from Northwestern’s Office of Undergraduate Research. RWC also acknowledges support from the National Science Foundation (1764421, 2235451) and Simons Foundation (597491, MP-TMPS-00005320).

---

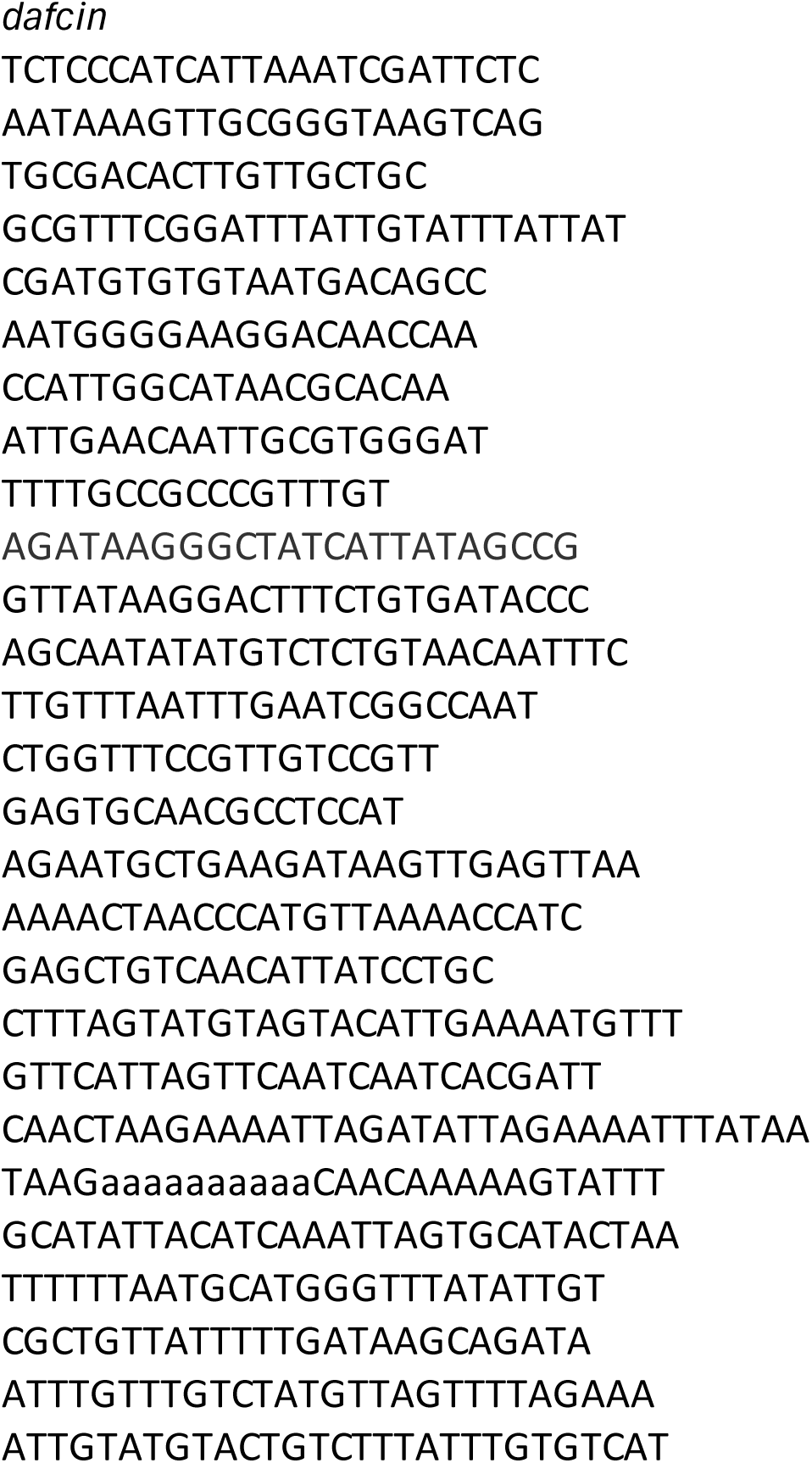

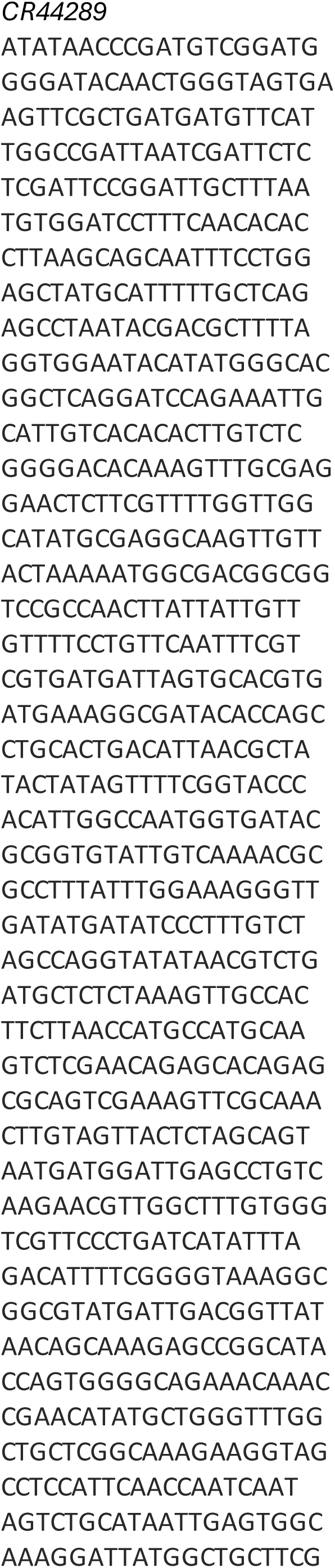
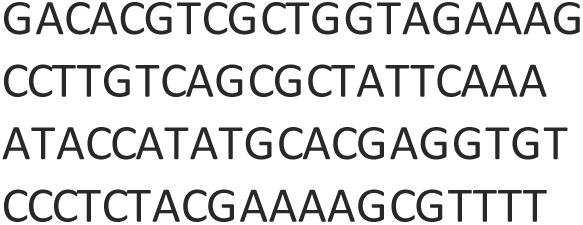

| **Species** | **Accession number** | **Source** | **tissue** |
| --- | --- | --- | --- |
| melanogaster | SRR070431 | SRA | ovaries |
| melanogaster | SRR100283 | SRA | ovaries |
| melanogaster | SRR7243502 | SRA | ovaries |
| melanogaster | SRR7243936 | SRA | ovaries |
| melanogaster | SRR7243550 | SRA | ovaries |
| melanogaster | SRR7243552 | SRA | ovaries |
| simulans | SRR618931 | SRA | ovaries |
| simulans | SRR2644782 | SRA | ovaries |
| simulans | SRR2644790 | SRA | ovaries |
| simulans | SRR2568366 | SRA | ovaries |
| simulans | SRR2568367 | SRA | ovaries |
| yakuba | SRR5639533 | SRA | ovaries |
| yakuba | SRR5639534 | SRA | ovaries |
| yakuba | SRR5639535 | SRA | ovaries |
| yakuba | SRR6181102 | SRA | ovaries |
| ananassae | SRR5639281 | SRA | ovaries |
| ananassae | SRR5639282 | SRA | ovaries |
| ananassae | SRR5639283 | SRA | ovaries |
| ananassae | SRR6181018 | SRA | ovaries |
| pseudoobscura | SRR5639407 | SRA | ovaries |
| pseudoobscura | SRR5639408 | SRA | ovaries |
| pseudoobscura | SRR5639409 | SRA | ovaries |
| pseudoobscura | SRR618060 | SRA | ovaries |
| pseudoobscura | DRR055260 | NIG | ovaries |
| pseudoobscura | DRR055261 | NIG | ovaries |
| willistoni | SRR5639491 | SRA | ovaries |
| willistoni | SRR5639492 | SRA | ovaries |
| willistoni | SRR5639493 | SRA | ovaries |
| willistoni | SRR6181088 | SRA | ovaries |
| virilis | SRR5639449 | SRA | ovaries |
| virilis | SRR5639450 | SRA | ovaries |
| virilis | SRR5639451 | SRA | ovaries |
| virilis | SRR6181074 | SRA | ovaries |
| mojavensis | SRR5639323 | SRA | ovaries |
| mojavensis | SRR5639324 | SRA | ovaries |
| mojavensis | SRR5639325 | SRA | ovaries |
| grimshawi | SRR3355200 | SRA | ovaries |
| grimshawi | SRR3355201 | SRA | ovaries |
| grimshawi | SRR3355202 | SRA | ovaries |
| grimshawi | SRR3355203 | SRA | ovaries |
| grimshawi | SRR3355262 | SRA | ovaries |
| grimshawi | SRR3355263 | SRA | ovaries |
| grimshawi | SRR3355264 | SRA | ovaries |
| grimshawi | SRR3355265 | SRA | ovaries |

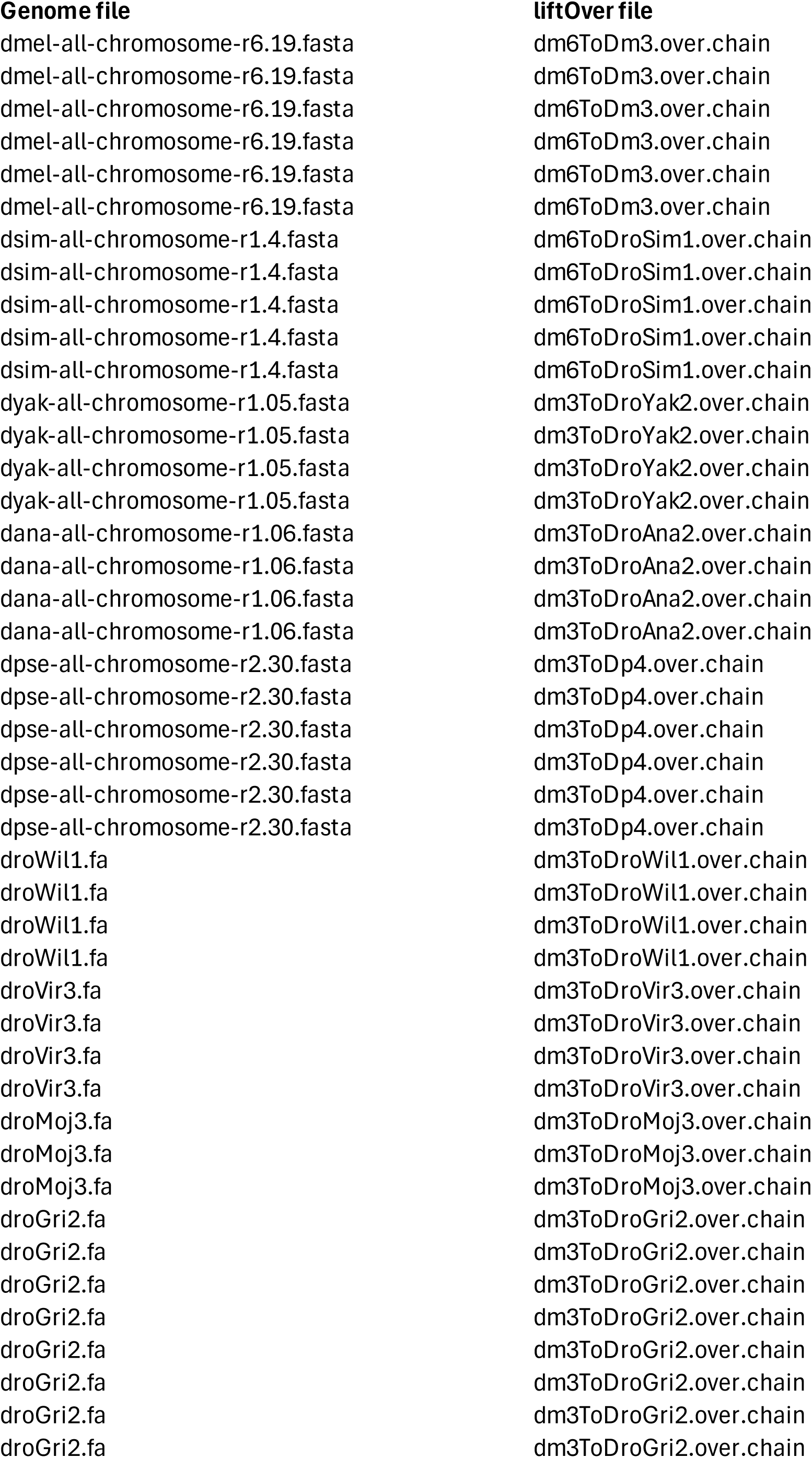

## Notes

### Competing Interest Statement

The authors have declared no competing interest.

